# Integrated lipidomics and proteomics network analysis highlights lipid and immunity pathways associated with Alzheimer’s disease

**DOI:** 10.1101/2020.03.18.995464

**Authors:** Jin Xu, Giulia Bankov, Min Kim, Asger Wretlind, Jodie Lord, Rebecca Green, Angela Hodges, Abdul Hye, Dag Aarsland, Latha Velayudhan, Richard J.B. Dobson, Petroula Proitsi, Cristina Legido-Quigley, on behalf of the AddNeuroMed Consortium

**Affiliations:** Institute of Pharmaceutical Science, King’s College London, United Kingdom; Institute of Psychiatry, Psychology and Neuroscience, King’s College London, United Kingdom; Steno Diabetes Centre, Copenhagen, Denmark

**Author notes:** Corresponding authors; E-mail address (R.J.B.D); E-mail address (P.P); E-mail address (C.L.Q).

**Keywords:** Alzheimer’s disease, Dementia, Brain atrophy, sMRI, Rate of cognitive decline, Lipidomics, Proteomics, AD risk loci, WGCNA

## Abstract

**INTRODUCTION:** There is an urgent need to understand the molecular mechanisms underlying Alzheimer’s Disease (AD) to enable early diagnosis and develop effective treatments. Here we aim to investigate Alzheimer’s dementia using an unsupervised lipid, protein and gene multi-omic integrative approach.

**METHODS:** A lipidomics dataset (185 AD, 40 MCI and 185 controls) and a proteomics dataset (201 AD patients, 104 MCI individuals and 97 controls) were utilised for weighted gene co-expression network analyses (WGCNA). An additional proteomics dataset (94 AD, 55 MCI and 100 controls) was included for external proteomics validation. Modules created within each modality were correlated with clinical AD diagnosis, brain atrophy measures and disease progression, as well as with each other. Gene Ontology (GO) enrichment analysis was employed to examine the biological processes and molecular and cellular functions for protein modules associated with AD phenotypes. Lipid species were annotated in the lipid modules associated with AD phenotypes. Associations between established AD risk loci and lipid/protein modules that showed high correlation with AD phenotypes were also explored.

**RESULTS:** Five of the 20 identified lipid modules and five of the 17 identified protein modules were correlated with AD phenotypes. Lipid modules comprising of phospholipids, triglycerides, sphingolipids and cholesterol esters, correlated with AD risk loci involved in immune response and lipid metabolism. Five protein modules involved in positive regulation of cytokine production, neutrophil mediated immunity, humoral immune responses were correlated with AD risk loci involved in immune and complement systems.

**DISCUSSION:** We have shown the first multi-omic study linking genes, proteins and lipids to study pathway dysregulation in AD. Results identified modules of tightly regulated lipids and proteins that were strongly associated with AD phenotypes and could be pathology drivers in lipid homeostasis and innate immunity.

**Research in Context:**

1. Lipid and protein modules were preserved amongst Alzheimer’s disease (AD) patients, participants with mild cognitive impairment (MCI) and controls. Protein modules were also externally validated.
2. Five lipid and five protein modules out of a total of thirty-seven correlated with clinical AD diagnosis, brain atrophy measurements and the rate of cognitive decline in AD.
3. Lipid and protein modules associated with AD phenotypes showed associations with established AD risk loci involved in lipid and immune pathways.

## Introduction

There is an urgent need to better understand the molecular mechanisms underlying Alzheimer’s Disease (AD) to enable early diagnosis and develop effective treatments. With the estimated worldwide number of patients suffering from dementia rising up to 115.4 million in 2050 [1], AD is undoubtedly one of the major healthcare challenges of the 21st century. Blood-based biomarkers could act as an easily accessible and minimally invasive screening tool to identify at risk individuals for further investigation and monitoring or stratification in clinical trials. Additionally they can reveal the molecular pathway leading to AD, creating new opportunities for drug development [2].

In the past decade a large number of untargeted and targeted blood biomarker studies have nominated and replicated proteins and combinations of proteins associated with AD and AD endophenotypes. These endophenotypes include brain atrophy, rate of cognitive decline and amyloid burden [3, 4]. Although the majority of protein biomarkers have failed to replicate, several proteins, especially inflammatory proteins and proteins involved in the complement pathway, have been consistently associated with AD or AD endophenotypes, including complement C6 and C-C motif chemokine 15 [3].

More recently, a number of untargeted and targeted blood metabolomic studies have emerged, highlighting the role of lipids in AD [5-7]. Lipidomics aims to identify and quantify thousands of lipids. It is regarded as a subset of metabolomics [8, 9], reflecting functional networks of downstream changes of the genome, transcriptome and proteome [10], and bridging the phenotype–genotype gap due to their close association to cellular processes [11]. We have previously performed lipid phenotyping and identified a panel of ten metabolites which predicted an AD training dataset with 83% accuracy and a test dataset with 79% accuracy [12]. As in the case of proteins, results have not always replicated, but Phosphatidylcholines (PCs), Cholesteryl Esters (ChEs), and Triglycerides (TGs) have been consistently shown alterations in MCI and AD compared to controls [7, 12-14].

Most biomarker studies to date have been restricted to one modality (proteomics or metabolomics), and only a modest number have used systems biology approaches [15-17]. Network analysis methods provide a powerful tool to depict higher order interactions between genes or proteins which can pinpoint disease-associated networks of highly connected molecules that represent important targets for understanding and treating AD. A small number of blood and brain network studies using Weighted Gene (or Protein or Lipid) Correlation Network Analysis (WGCNA) have highlighted gene, protein and lipid pathways that are involved in the aetiology, initiation, and progression of AD [15, 18].

Here, we aimed to explore the role of blood lipids and proteins in AD at a systems level by performing an integrative multiscale network analysis and correlating the identified modules with clinical AD diagnosis, brain atrophy and the rate of cognitive decline. Encouraged by genome-wide association studies (GWAS) and meta-analyses consistently implicating immunity and lipid processing in AD [19], we additionally integrated these networks with established AD risk loci. Our study highlights the use of large-scale integrated systems biology approaches to unravel the molecular aetiology of AD. The study design is illustrated in **Figure 1**.

**Figure 1.**
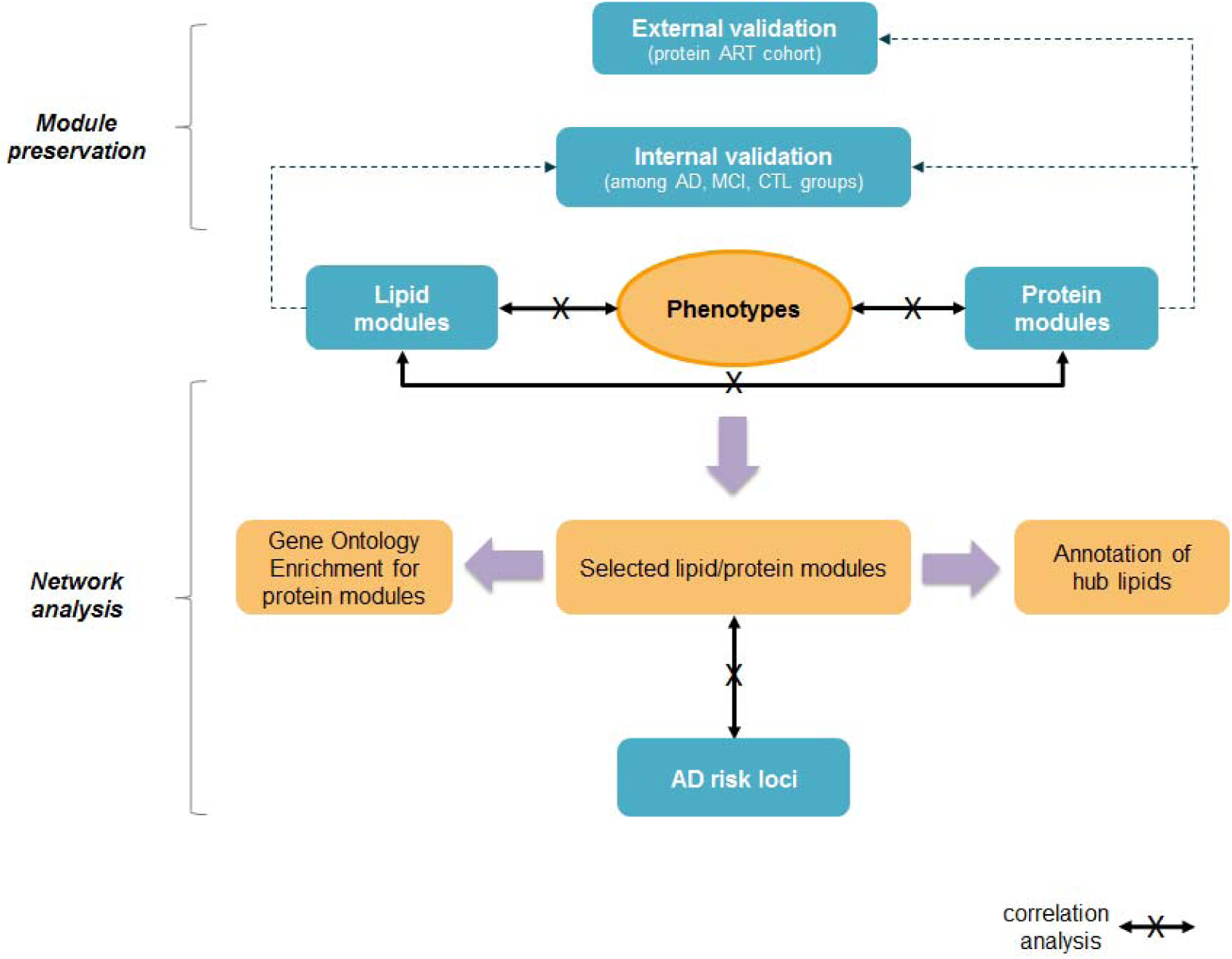
Study workflow. Protein and lipid modules were produced, and their preservation was investigated. An internal validation among AD, MCL and CTL groups were done for protein and lipid modules, additional external validation of the protein modules was performed against the ART cohort. Correlation analysis between lipid modules, protein modules and phenotypes (clinical diagnosis, rate of cognitive decline, hippocampal left and right volume, entorhinal cortex left and right volume) were made separately and led to a selected number of modules. Gene ontology enrichment analysis was applied for selected protein modules, while the annotation of lipid species for selected lipid modules was conducted. The association between lipid/protein modules and AD risk loci was investigated..

## Methods

### Subjects

This study included three datasets as listed in **Table 1**. Briefly, dataset 1 contained proteomic data across 201 AD patients, 104 individuals with mild cognitive impairment (MCI) and 97 controls from the EU-funded AddNeuroMed (ANM) study [20]. Dataset 2 consisted of lipidomic data across 185 AD, 40 MCI and 185 controls from the Maudsley and King’s Healthcare Partners Dementia Case Register (DCR) and the ANM study. The overlap between dataset 1 and 2 includes 240 individuals: 147 AD, 10 MCI and 83 controls. An additional proteomics dataset (dataset 3) from the Alzheimer’s Research Trust (ART) cohort [21] was included with 94 AD, 55 MCI and 100 controls.

**Table 1.** Sample demographics

| **Proteomics dataset**<br>**(dataset 1, N = 411)** | **AD**<br>**(N = 210)** | **MCI**<br>**(N = 104)** | **Controls**<br>**(N = 97)** | **Differences among**<br>**three groups\*** |
| --- | --- | --- | --- | --- |
| Age, mean (SD) | 77 (6.5) | 75 (6.1) | 72 (6.6) | $P = 1.13\text{e-}08$ |
| Gender (female/male) | 133/77 | 59/45 | 56/41 | $P = 0.4431$ |
| *APOE* ε4 allele<br>(absence/presence) | 89/115 | 58/46 | 65/32 | $P = 5.349\text{e-}04$ |
| ROD (per year),<br>mean (SD)<sup>a</sup> | - 1.49 (1.26)<br>(N = 127) | NA | NA | NA |
|  | **AD**<br>**(N = 53)** | **MCI**<br>**(N = 0)** | **Controls**<br>**(N = 67)** | **Differences among**<br>**three groups\*** |
| Whole brain volume,<br>Mean (SD)<sup>c</sup> | 0.66 (0.073) | NA | 0.69 (0.043) | $P = 0.003184$ |
| Hippocampus left,<br>mean (SD)<sup>c</sup> | 0.0018<br>(0.00037) | NA | 0.0024<br>(0.00031) | $P = 7.717\text{e-}15$ |
| Hippocampus right,<br>mean (SD)<sup>c</sup> | 0.0019<br>(0.00041) | NA | 0.0025 (0.00031) | $P = 3.535\text{e-}13$ |
| Entorhinal cortex left,<br>mean (SD)<sup>c</sup> | 0.00095<br>(0.00038) | NA | 0.0012 (0.00024) | $P = 1.433\text{e-}05$ |
| Entorhinal cortex right,<br>mean (SD)<sup>c</sup> | 0.00092<br>(0.00034) | NA | 0.0013 (0.00028) | $P = 3.562\text{e-}08$ |
| **Lipidomics dataset**<br>**( dataset 2, N = 415)** | **AD**<br>**(N = 185)** | **MCI**<br>**(N = 40)** | **Controls**<br>**(N = 190)** | **Differences among**<br>**three groups\*** |
| Age, mean (SD) | 77 (6.9) | 75 (6.3) | 79 (5.5) | $P = 6.77\text{e-}05$ |
| Gender (female/male) | 114/71 | 20/20 | 116/74 | $P = 0.377$ |
| *APOE* ε4 allele<br>(absence/presence) | 73/110 | 22/15 | 138/51 | $P = 8.725\text{e-}10$ |
| ROD (per year),<br>mean (SD)<sup>a</sup> | -1.50 (1.22)<br>(N = 144) | NA | NA | NA |
|  | **AD**<br>**(N = 57)** | **MCI**<br>**(N = 8)** | **Controls**<br>**(N = 67)** | **Differences among**<br>**three groups\*** |
| Whole brain volume,<br>Mean (SD)<sup>c</sup> | 0.66 (0.071) | 0.71 (0.041) | 0.69 (0.098) | $P = 0.0006595$ |
| Hippocampus left,<br>mean (SD)<sup>c</sup> | 0.0018<br>(0.00038) | 0.0021<br>(0.00031) | 0.0024 (0.00060) | $P = 1.763\text{e-}15$ |
| Hippocampus right,<br>mean (SD)<sup>c</sup> | 0.0018<br>(0.00043) | 0.0022<br>(0.00032) | 0.0024 (0.00062) | $P = 2.572\text{e-}13$ |
| Entorhinal cortex left,<br>mean (SD)<sup>c</sup> | 0.00093<br>(0.00038) | 0.0012<br>(0.00024) | 0.0012 (0.00030) | $P = 2.782\text{e-}06$ |
| Entorhinal cortex right,<br>mean (SD)<sup>c</sup> | 0.00090<br>(0.00033) | 0.0011<br>(0.00028) | 0.0013 (0.00027) | $P = 4.114\text{e-}09$ |
| **Proteomics validation**<br>**dataset**<br>**( dataset 3, N = 249)** | **AD**<br>**(N = 94)** | **MCI**<br>**(N = 55)** | **Controls**<br>**(N = 100)** | **Differences among**<br>**three groups\*** |
| Age, mean (SD) | 83 (6.2) | 77 (4.5) | 79 (7.0) | $P = 2.02\text{e-}12$ |
| Gender (female/male) | 72/22 | 35/20 | 47/53 | $P = 1.185\text{e-}04$ |
| *APOE* ε4 allele<br>(absence/presence) | 40/50 | 30/10 | 79/20 | $P = 8.204\text{e-}07$ |

AD, Alzheimer's disease; ROD, rate of cognitive decline; SD, standard deviation; NA, not available
\*Differences in the means/frequencies of clinical/demographic variables were tested using ANOVA, t test or x2 test.
<sup>a</sup> Rate of cognitive decline data was available for a subset of AD patients (N1 = 127, N2 = 144).
<sup>c</sup> sMRI data were available for a subset of study participants (N1 = 120 [AD = 53, controls = 67], N2 = 132 [AD = 57, MCI = 8, controls = 67]).

All individuals with AD met criteria for either probable (NINCDS-ADRDA, DSM-IV) or definite (CERAD) AD. Individuals in the control group were screened for dementia using the mini mental state examination (MMSE) or ADAS-cog. They were determined to be free from dementia at neuropathologic examination or had a Braak score. Diagnosis was confirmed by pathologic examination for a proportion of cases and cognitively normal elderly controls. All AD cases had an age of onset of at least 60 years, and controls were 60 years or above at examination. Each individual was required to fast for 2 hours before sample collection, and 10 mL of blood was collected in tubes coated with sodium ethylenediaminetetraacetic acid to prevent clotting. Whole blood was centrifuged at 2000 g for 10 minutes at 4°C to separate plasma, which was removed and stored at - 80°C. All samples were centrifuged within approximately 2 hours of collection.

### Proteomics and lipidomics analyses

The details of lipidomics and proteomics experiments are described elsewhere [14, 22]. Overall, 1016 proteins with Uniprot ID were measured with a Slow Off-rate Modified Aptamer (SOMAmer)–based capture array called “SOMAscan” (SomaLogic, Inc). Lipidomics was performed by a Waters ACQUITY UPLC and XEVO QTOF system where 2216 lipid features were measured and included.

The detailed methods of structural magnetic resonance imaging and calculation of rate of cognitive decline can be found in the supplementary document, where the descriptions about weighted lipid co-expression network and weighted protein co-expression network are also included.

### Associations between lipid and protein modules

The associations between five lipid and five protein modules associated with AD phenotypes at a Bonferroni corrected level were also investigated with Pearson’s correlation. The correlations of individual lipids to proteins in modules that passed Bonferroni correction (*p* = 2e-03) were further investigated at two different thresholds with correlation coefficient absolute values in the range of 0.1 to 1 and 0.2 to 1.

### Annotation of top lipid module drivers and gene set enrichment analyses of protein modules

Lipid species were annotated in the selected lipid modules i.e. those associated with at least two phenotypes. To examine the biological processes and the molecular and cellular functions, Gene Ontology (GO) enrichment analysis against KEGG and Reactome pathways and Over-representation analysis (ORA) were performed for all five protein modules associated with the tested phenotypes using WebGestalt (WEB-based Gene SeT AnaLysis Toolkit) [23]. The genome database was used as the background/reference set. Z scores determined the over-representation of ontologies in a module and one tailed Fisher’s exact tests (Benjamini-Hochberg FDR corrected) were used to assess the significance of the Z score [24]. A minimum of five genes per ontology were used as filters prior to pruning the ontologies and the FDR significant (q<0.05) or top 10 categories were selected.

### Lipid and protein modules association with AD genetic variants

Linear regression was used to investigate associations of 36 established AD risk variants [19, 25, 26] with the five lipid and five protein modules associated with the six tested phenotypes. A Bonferroni corrected p-value threshold of p<1.5e-04 was applied, however, p value less than 0.05 was considered to be significant.

## Results

Sample demographics are displayed in **Table 1**.

### Module preservation

To assess the reliability and reproducibility of the established modules and to investigate whether the modules were preserved between AD patients, individuals with MCI and healthy controls, module preservation analysis was applied. Most of the lipid and proteins modules showed medium-high preservation between AD cases, individuals with MCI and controls. Therefore, WGCNA was applied to all three groups (**Supplement Figures S1 and S2 respectively**). External validation was conducted for the ANM proteomics dataset by utilising the ART cohort as the test set/reference set (**Supplement Figure S3)**. Overall, there was good agreement between the ANM and ART cohort module assignments. Five ANM modules (black, blue, grey60, red and lightyellow) were well preserved in ART (black, green yellow, turquoise, blue and lightgreen respectively). Conversely, the ANM module midnightblue appeared not to be preserved in ART networks since most of its proteins were classified as unassigned (grey colour). The rest of the modules in the ANM network appeared to show medium preservation and split into one or more modules in the ART network. For example, lightcyan module in the ANM network showed high preservation with salmon and lightyellow modules in the ART network.

Weighted lipid correlation network analysis identified 17 modules of co-regulated lipids ranging from 36 to 328 lipids in each module (**Supplement Figure S4A**). Weighted protein correlation network analysis identified 17 modules of co-regulated proteins ranging from 20 to 159 proteins in each module (**Supplement Figure S4B**).

### Association of lipid modules with AD phenotypes

After Bonferroni correction, 11 lipid modules were associated with at least one trait (**Figure 2A)**. Most associations were observed with ROD, where five modules showed positive associations and three modules showed negative associations. Additionally, one module (green) was reduced in AD versus controls and five modules showed associations with brain atrophy, of which two were associated with less brain atrophy and three were associated with increased brain atrophy. Only three of the eight modules correlated with ROD were also associated with AD diagnosis or brain atrophy. Overall, there were five modules associated with at least two phenotypes. These included the darkturquoise module, which associated with faster ROD and greater atrophy; green module, which was decreased in AD and was also associated with less atrophy; greenyellow module, which was associated with greater atrophy; midnightblue module, which was associated with slower ROD and less atrophy in the hippocampus; and orange module, which was associated with greater brain atrophy and was increased in AD, though the association with AD did not survive multiple testing (**Figure 2B-F)**.

**Figure 2.**
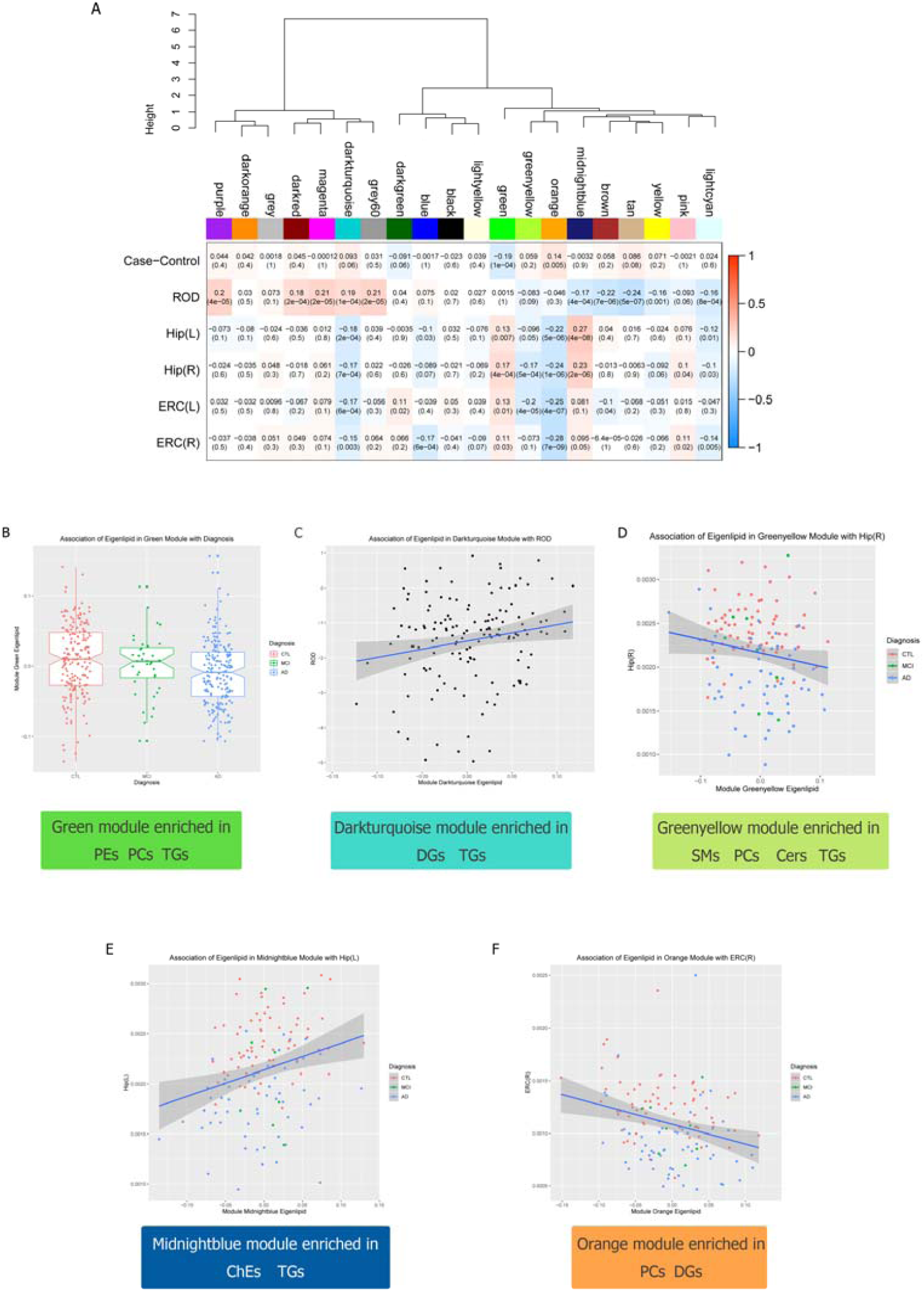
Correlations between lipid modules and AD phenotypes. (A) Lipid modules were clustered to assess module relatedness based on correlation of lipid co-expression eigenlipids (top). Heat map showing the correlation of lipid module eigenlipids and phenotypes (bottom). (B) Association of eigenlipid with diagnosis and main lipid species in green module. (C) Association of eigenlipid with ROD and main lipid species in darkturquoise module. (D) Association of eigenlipid with entorhinal cortex left volume and main lipid species in greenyellow module. (E) Association of eigenlipid with Hippocampal left volume and main lipid species in midnightblue module. (F) Association of eigenlipid with entorhinal cortex right volume and main lipid species in orange module.

We decided to further investigate the five modules associated with at least two phenotypes. The top drivers for each of the five modules based on their kME are listed in **Supplement Table S2.** Annotations for all lipids in each module revealed that the green module was enriched in phosphatidylethanolamines(PEs), phosphatidylcholines (PCs) and triglycerides (TGs); the orange module was enriched in PCs and diacylglycerides (DGs); the greenyellow module was enriched in sphingomyelins (SMs), PCs, Ceramides (Cers) and TGs; the darkturquoise module was enriched in TGs and DGs and the midnightblue module was enriched in TGs and ChEs (**Table S1**). We subsequently observed strong associations between the module membership of the lipids in the five modules and the lipids-phenotype correlations, indicating that lipids with high kME (i.e. top module drivers/ hub lipids) were also associated with the respective AD phenotypes. **Figure 2B** displays the association of the green module with clinical diagnosis and **Figure S9A** shows the association of lipid kME in the green module against the lipid-diagnosis correlation. The rest of the associations between lipid modules and phenotypes that passed Bonferroni correction are presented in Supplement **Figure S6** and **Figure S7**. The associations of lipid module memberships in the five selected modules and the lipid-phenotype correlations that passed Bonferroni correction are presented in Supplement **Figure S10**.

### Association of protein modules with AD phenotypes

After Bonferroni correction, five protein modules were associated with at least one trait (**Figure 3A)**. Four of the five modules (yellow, red, cyan and lightcyan) were associated with increased brain atrophy and one module (lightgreen) was increased in AD patients compared to controls. Most associations were observed for Hippocampus left volume and although some modules were associated with more than one phenotype, most of these associations did not pass multiple testing corrections (**Figure 3B-F**).

**Figure 3.**
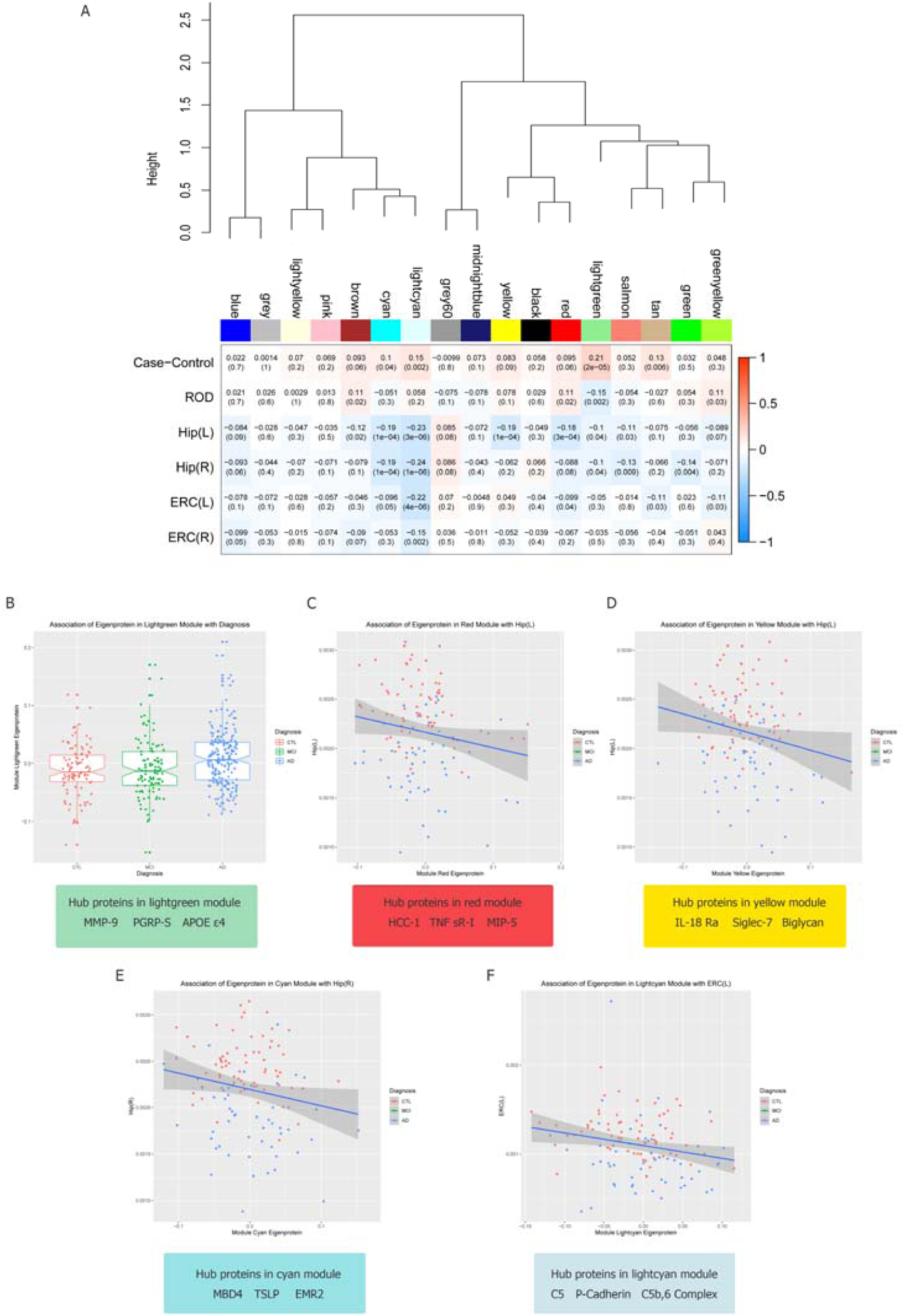
Correlations between protein modules and AD phenotypes. (A) Protein modules were clustered to assess module relatedness based on correlation of protein co-expression eigenprotein (top). Heat map showing the correlation between protein module eigenproteins and phenotypes (bottom). (B) Association of eigenprotein with diagnosis and hub proteins in lightgreen module. (C) Association of eigenprotein with hippocampal left volume and hub proteins in red module. (D) Association of eigenprotein with hippocampal left volume and hub proteins in yellow module. (E) Association of eigenprotein with hippocampal right volume and hub proteins in cyan module. (F) Association of eigenprotein with entorhinal cortex left volume and hub proteins in lightcyan module.

The top drivers for each of the five protein modules, based on their kME, are displayed in **Supplement Table S3**. Overall, it was observed that module membership for proteins was lower compared to that of lipids. The top drivers in the five modules included many AD protein candidates, some of which have been associated with AD in the same cohort. ApoE, one of these top drivers, is involved in complement cascade and growth factors. GO over-representation analyses results for proteins in the five modules associated with AD phenotypes are presented in (**Supplement Table S4**). The biological processes that passed the Benjamini-Hochberg (BH) correction for each protein module are shown in the directed acyclic graph (DAG) in **Figure S5**. Biological GO terms that passed BH correction for the lightgreen module included neutrophil mediated immunity, granulocyte activation, STAT cascade and regulation of inflammatory response; biological GO terms for the cyan module included positive regulation of cytokine production and mast cell activation; biological GO terms for the lightcyan module included protein activation cascade, insulin-like growth factor receptor signaling pathway and regulation of plasma lipoprotein particle levels; and biological GO terms for the red and yellow modules included humoral immune response. For KEGG pathway analysis, GO terms like complement and coagulation cascades which passed BH correction in lightcyan and yellow module, was also included in cyan module; JAK-STAT signaling pathway was listed as one of the top 10 pathways in lightgreen module and cyan module; Mapk signaling pathway was highlighted in red module after BH correction.

When we plotted kME for the proteins in the five modules versus the proteins-phenotype correlations, with the exception of the lightgreen module and AD diagnosis, correlations were not as strong as in the case of lipids. **Figure 3B** displays the association of the lightgreen module with clinical diagnosis and **Figure S9B** illustrates the association of protein kME in the lightgreen module against the protein-diagnosis correlation. The rest of the associations between protein modules and phenotypes that passed Bonferroni correction are presented in Supplement **Figure S8**. The associations of protein module memberships against the protein-phenotype correlations that passed Bonferroni correction are presented in Supplement **Figure S11**.

### Association between lipid modules and protein modules

The relationship between lipid modules and protein modules associated with AD phenotypes was also investigated. As shown in **Figure 4A**, the greenyellow lipid module was positively correlated with the lightcyan protein module after correcting for multiple testing. Additionally, the darkturquoise lipid module was positively correlated with the lightgreen protein module. The summary of associations among five lipid modules, five protein modules and six phenotypes are illustrated by the circus plot in **Figure 4B**. The correlations of individual lipids in the greenyellow module and proteins in the lightcyan module, as well as lipids in the darkturquoise module and proteins in the lightgreen module were further investigated at two different thresholds (Supplement **Figure S12** and **Figure S13**). Strong correlations were observed between ApoB, PAFAH, P-cadherin in the lightcyan module and a group of lipids including some top drivers in the greenyellow module and DG (38:2). Similarly, ApoE ε3 and ApoE ε4 in the lightgreen module showed strong correlation with top lipid drivers in the darkturquoise module.

**Figure 4.**
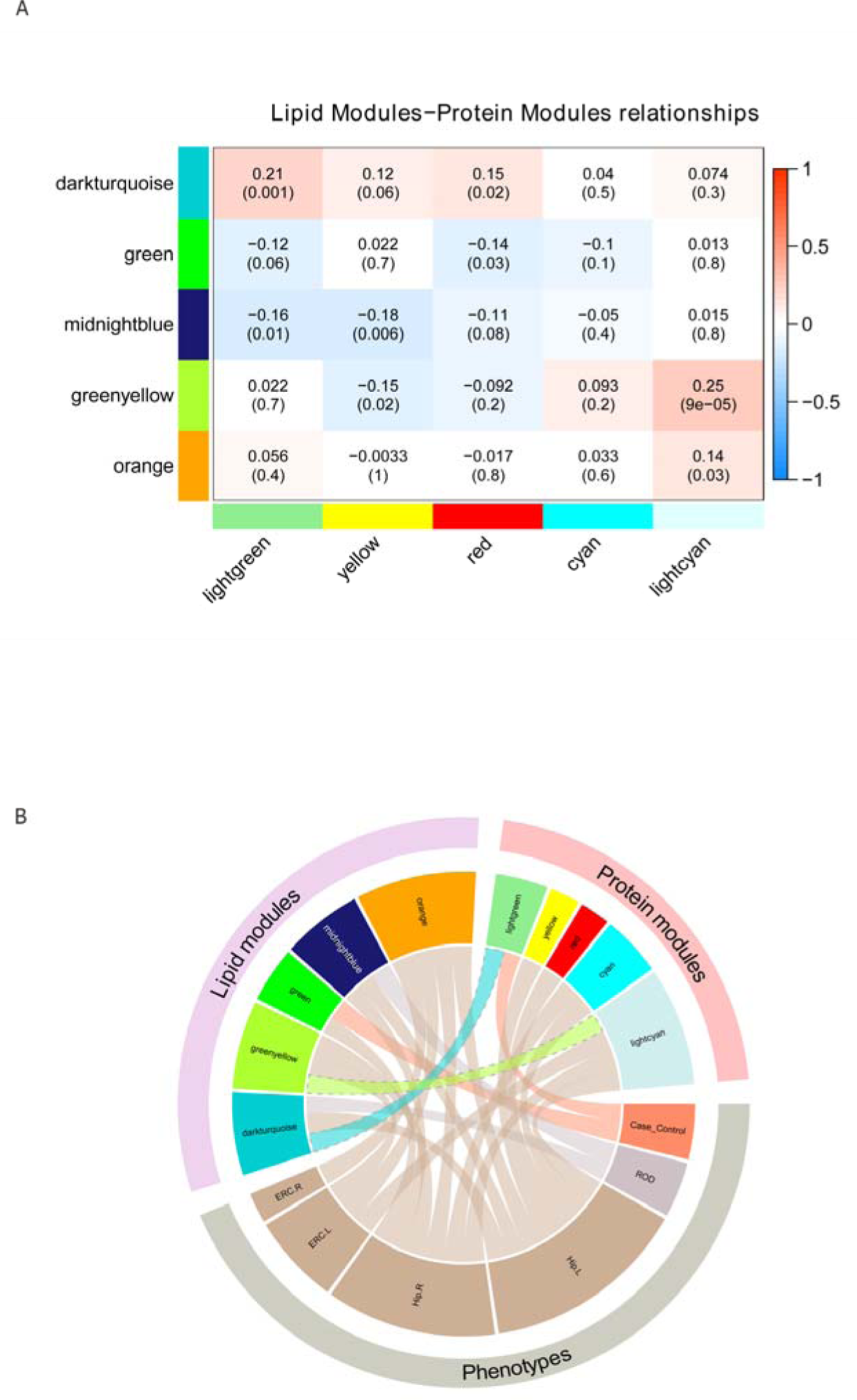
Associations between co-expression protein networks and lipid networks and phenotypes. (A) Heat map showing the Pearson correlations and p values (in bracket) between 5 lipid modules (rows) associated with phenotypes and 5 protein modules (columns) associated with phenotypes; (B). Circos Plot showing the correlation among 5 lipid modules, 5 protein modules and 6 phenotypes.

### Association between AD genetic variants and selected lipid modules/protein modules

The five lipid modules associated with more than one AD phenotype (darkturquoise, green, greenyellow, midnightblue and orange modules), and all five protein modules associated with AD phenotypes (lightgreen, red, yellow, cyan and lightcyan modules) were selected for further analysis with 36 AD genetic variants [19, 25, 26].

Three of the four lipid modules showed association with six genetic variants, while four out of five protein modules demonstrated association with nine genetic variants at *p*< 0.05. However, none of these associations passed correction for multiple testing (listed in **Table S5)**. The summarised association for lipid modules and AD risk loci, as well as protein modules and AD risk loci are shown in **Figure 5**.

**Figure 5.**
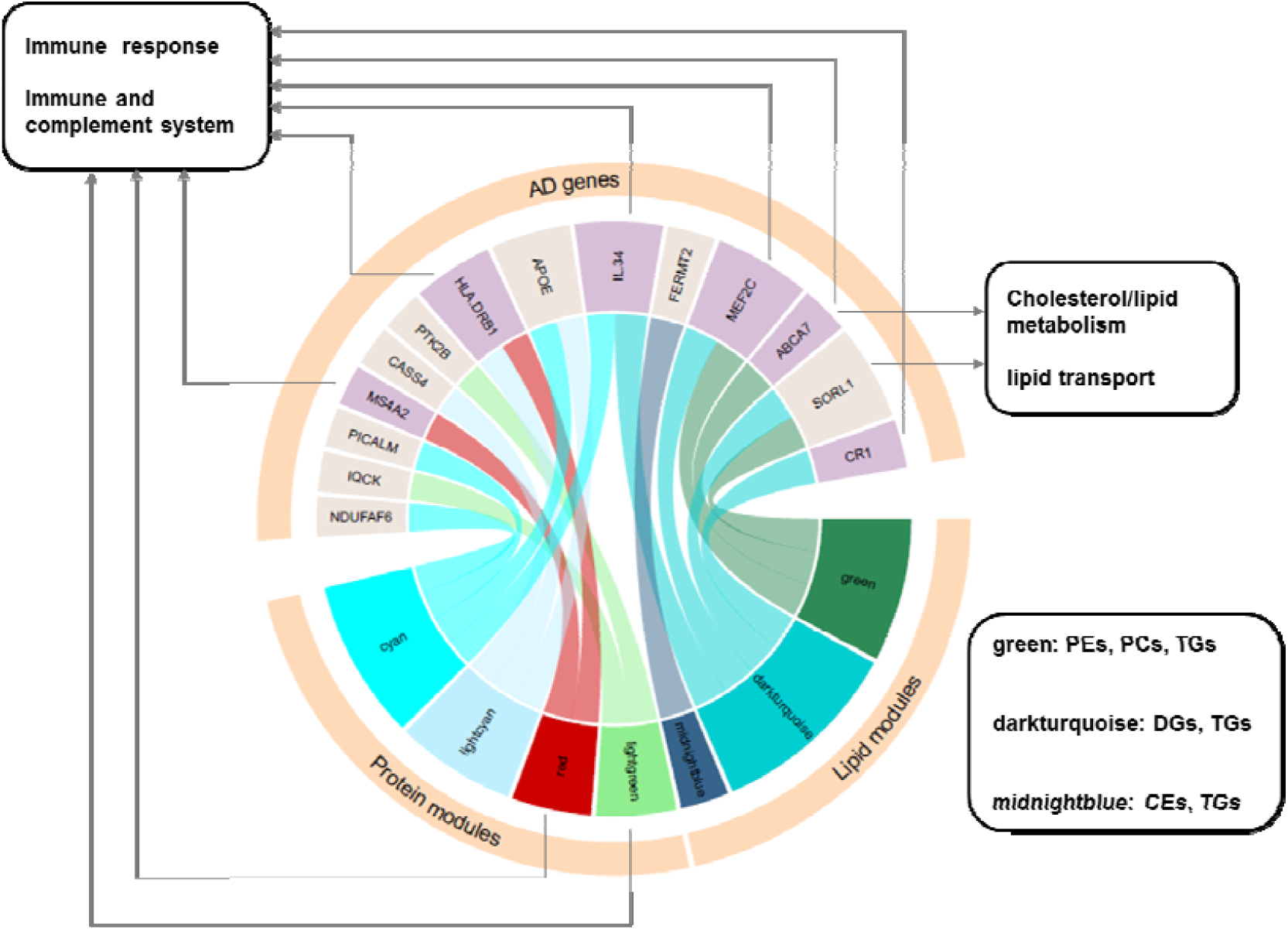
Summary of lipid modules/protein modules association with AD risk loci. AD genetic variants IL34, MEF2C, ABCA7, SORL1 and CR1, correlated with lipid module green, darkturquoise, and midnightblue, has established functions of immune response and inflammation, cholesterol metabolism, lipid transport and immune and complements systems. Two protein modules, cyan and red, were linked with immune response through over-representation analysis, while AD genetic variants MS4A2, HLA-DLB1 and IL34, correlated with protein modules, also showed functions of immune response.

None of the lipid modules were associated with the ApoE ε4 allele. Presence of the ApoE ε4 allele was associated with increased levels of the protein modules cyan and lightcyan (*p*=0.0048 and *p*=0.0013 respectively).

## Discussion

In this study, we employed a multi-omic network approach on data available for a total of 586 participants: 415 with lipidomics data, 411 with proteomics data, and 249 with proteomics validation data from three prospective cohorts. Overall, we identified modules of tightly regulated lipids and proteins that were strongly associated with AD phenotypes. Associations were more abundant for lipid modules, with the majority of these with ROD, followed by brain volume. Conversely, most protein modules demonstrated associations with brain atrophy, particularly with hippocampal volume. A number of lipid and protein clusters were also observed to be correlated with each other. Finally, AD associated modules demonstrated modest associations with AD genes involved in lipid and immune processes. Individual lipid/protein were grouped using network analysis and associated with relevant clinical parameters [27].

Lipid co-expression network analysis revealed strong correlations between 11 lipid modules and AD diagnosis, ROD and brain atrophy measurements. These modules were preserved amongst AD, MCI and control groups; thus, the clustering of lipid features was not affected by clinical diagnosis. Not all modules were associated across the spectrum of tested phenotypes with some being restricted to brain atrophy or ROD, suggesting distinct biological mechanisms. In addition, the five lipid modules associated with more than one phenotype were enriched in PCs, TGs and ChEs, which have shown to be linked with AD in our studies including analyses on this cohort [12-14]. For example, the green module, which is enriched in TGs, is reduced in MCI and healthy control groups compared to AD cases, and positively correlated with entorhinal and hippocampal volume. We also observed that a lipid feature annotated as TG (60:11), with a mass to charge ratio of 970.79, was previously associated with AD diagnosis and hippocampal volume using machine learning on overlapping samples [14]. This was one of the top drivers in the lipid green module, highlighting the importance of big TGs as possible drivers in AD processes. A recent study using the Alzheimer’s Disease Neuroimaging Initiative (ADNI) cohort highlighted the association of TGs and especially poly-unsaturated fatty acid containing TGs with AD and AD biomarkers [28]. TGs are also associated with Aβ, and longitudinal studies have shown that midlife TGs are associated with amyloidosis and tau pathology 20 years later in cognitively healthy individuals [29].

The protein modules identified were highly preserved through internal validation, as well as in an independent proteomic cohort. Five of these modules were strongly associated with AD phenotypes. With the development of analytical platforms, increasing numbers of studies have attempted to find AD biomarkers in blood. From the nominated biomarkers, only some have been replicated. The reason for such imbalance might be caused by the heterogeneity of the disease itself as well as the complexity of analysing blood [3]. Meta-analyses will help translate discovery findings to reproducible and useful biomarkers. In this study, we observed that some of the top drivers of the five AD associated modules (**Table S3**) were linked to AD pathology in regression and random forest analyses on the same cohort; these include C-C motif chemokine from the red module which associated with left hippocampus atrophy; carbonic anhydrase III and peptidoglycan recognition protein 1 from the lightgreen module which correlated with AD diagnosis [22]. Additionally, matrix metallopeptidase 9 [30, 31] and ApoE [31-35] from the lightgreen module, together with complement component C6 [36] from the lightcyan module were previously highlighted in other AD biomarker studies. These findings underline the importance of these proteins as drivers of networks and pathways associated with AD.

Over-representation analysis of the five AD related protein modules highlighted immune responses as the main biological process. KEGG pathways also highlighted the involvement of the complement, cytokine, MAPK signalling and the JAK-STAK pathways. Further, we observed correlation between lipid and protein modules associated with AD phenotypes, such as the association between the lightcyan protein module (enriched in complement proteins) and the greenyellow lipid module, suggesting important relationships and pathways that warrant further investigation. One of the GO biological terms revealed for the lightcyan protein module was the regulation of plasma lipoprotein particle levels. Additionally, molecular and cellular processes also highlighted protein-lipid complex and phospholipid binding and KEGG pathways including glycophospholipid biosynthesis (although these were associated at FDR <0.1).

Finally, when we investigated the association of the AD related lipid and protein modules we observed that AD risk loci CR1, SORL1, MEF2C, IL34 and ABCA7 were associated with several AD related lipid modules. These loci were involved in immune and complement system [37], lipid transport [38], immune response and inflammation [38, 39], cytokine signalling in immune system [40], as well as cholesterol/lipid metabolism, immune and complement systems [41], respectively. We also observed that protein modules which highlighted immune response as the main biological process were nominally associated with genetic variation at the IL34, HLA-DRB1 and MS4A (gene cluster) risk loci, which are all related to immune response, immune and complement systems [38, 41, 42]. Although none of these associations passed correction for multiple testing, this is likely due to the small sample size of the study and therefore these findings merit further investigation.

A limitation of our study is that the sample size was small. Further replication is therefore warranted, especially for genetic association analyses. Moreover, although we had overlapping lipidomics and proteomics data, these were available for only a subset of the study participants. Additionally, although we had information for ROD, this calculation was based on the MMSE, which is only one measure of cognition.

Nevertheless, our study highlights that the interpretation of multi-omic data such as lipidomic, proteomic and genomic can be boosted by deploying systems biology approaches. Our integrative approach highlights tightly regulated and inter-connected networks of lipids and proteins associated with AD and AD phenotypes with lipid and immunity pathways at its centre.

## Notes

#### Summary of Updates

Corresponding authors updated

