## Supplemental methods, figures and tables for "Integrated lipidomics and proteomics network analysis highlights lipid and immunity pathways associated with Alzheimer’s disease"

*Corresponding authors:

**Structural magnetic resonance imaging**

Volumes of whole brain (WB) and the hippocampi (HIP) and entorhinal cortices (ERC) were obtained from subjects who had undergone structure magnetic resonance imaging (sMRI) (**Table 1**). Regions were normalized by intracranial volume (1). Detailed information regarding data acquisition, pre-processing, and quality control assessment are described elsewhere (2, 3). Before analyses, sMRI measures were standardized to have a mean of 0 and a standard deviation (SD) of 1.

**Calculation of rate of cognitive decline**

The rate of cognitive decline (ROD) was available for 127 AD patients in dataset 1 and 144 AD patients in dataset 2. The ROD was calculated based on longitudinal MMSE assessments (4), and only samples with at least three MMSE measures were included for linear mixed effect models (5). Calculations were done separately for ANM and DCR due to the differences in assessment windows between the cohorts. After covariate adjustment, the slope coefficient obtained from the final model for each sample was then used as the ROD, defined as the change in MMSE per day (5).

**Weighted Lipid Co-expression Network Analysis**

Prior to network analysis, missing values were imputed using the k-nearest neighbour (KNN) imputation function from “impute” package within R version 3.6.1 (6). Inverse normal transformation (quantile normalisation) was applied to normalise the lipidomics data. Principal component analysis (PCA) was conducted on the genotype data to extract principal components that represent the population structure. Each normalised lipid was then regressed against age, gender, batch and the first seven principal components. The lipid residuals were used in downstream analyses.

A weighted lipid co-expression network was built with R package “WGCNA” (7) using the inverse-normalized lipidomics residuals. Briefly, a thresholding power of 12 was chosen (as it was the smallest threshold that resulted in a scale-free R2 fit of 0.9) and the signed network was created by calculating the component-wise minimum values for topologic overlap (TO). Using 1 – TO (dissTOM) as the distance measure, lipid features were hierarchically clustered. Initial module assignments were determined by using a dynamic tree-cutting algorithm (“tree” method, cutHeight=0.99, deepSplit = True, minModulesize = 30).

Before applying the network analysis to the whole lipidomics dataset, internal module preservation was applied to verify that module assignments were not affected by diagnosis groups. This was conducted by splitting lipidomics dataset into three sub datasets according to clinical status (AD, MCI, control). Module preservation analysis was applied for any two sub datasets, assigning one as the reference dataset and the other as the test dataset.

The resulting 20 modules or clusters of co-expressed lipid features were selected by merging modules based on the clustering of module eigenlipids (MEs; or the 1st principal component of the module). Module membership (kME) quantifies how close a lipid is to a given module and can be measured by calculating the correlation between individual lipids and the ME. Lipids with high kME (top 10) in the module were informally referred to as top drivers (8).

Association between the eigenlipids of 20 modules and six phenotypes including AD clinical diagnosis (AD vs Controls), hippocampal left and right volume, entorhinal cortex left and right volume and the ROD, were investigated using Pearson correlation. Only modules associated with more than one phenotype and with at least one passing Bonferroni correction threshold (*p* = 4.2e-04) were selected for further analysis. The relationship between lipid correlation with AD phenotypes and lipid module membership in selected modules was also investigated to examine the association of top module drivers with the examined phenotypes.

**Weighted Protein Co-expression Network Analysis**

Similar pre-treatment approaches were applied to the proteomics datasets. After imputing missing values using KNN (6), log2 transformation was applied and values outside of 4 standard deviations of the mean were excluded. The log2 transformed and outlier removed proteomics data were regressed against age, gender and the first seven PCs (for ANM cohort). The resulting datasets were utilised for further analyses.

Two weighted protein co-expression networks were built for the above pre-treated datasets using WGCNA (7). A thresholding power of 7 was chosen and the signed networks for both datasets were created following the same steps described in the weighted lipid co-expression network section, using a minimum module size of 17 (9). For both proteomics datasets, 17 modules were built.

Internal preservation analyses as previously described were applied in each proteomics dataset to investigate whether the generated modules were preserved in AD patients, controls and MCI individuals. External validation was also conducted using dataset 1 as reference, dataset 3 as test and vice versa. Cross-tabulation analysis was then employed to investigate module overlap and correlations between the two datasets (10). After validating the modules, Pearson’s correlation with Bonferroni correction (*p*=4.9e-04) was applied to investigate the association between the six phenotypes and the module eigenproteins for 17 modules in dataset 1. The relationship between individual protein correlations with AD phenotypes and protein module membership kME in selected modules was also explored as in the case of lipid modules.

**Table S1. Selected lipid modules and annotation.**

| Module | Association with phenotypes | Lipid species |
| --- | --- | --- |
| green | Case-control, Hip (L) | PE, PC, TG |
| darkturquoise | ROD, Hip(L) | DG, TG |
| midnightblue | ROD, Hip(L), Hip(R) | ChE, TG |
| greenyellow | Hip(R), ERC(L) | SMs, PCs, Cers, TGs |
| orange | Hip(L), Hip(R), ERC(L), ERC(R) | PC, DG |

PE, phosphatidylethanolamine; PC, phosphatidylcholine; TG, triglyceride; DG, diacylglyceride; CE, cholesteryl ester; SM, sphingomyelin; Cer, Ceramide.

**Table S2. Top 10 drivers in five selected lipid modules.**

| Module | Lipid feature/Putative annotation | kME |
| --- | --- | --- |
| darkturquoise | 604.54_29.19 | 0.93 |
|  | 603.54_29.19 | 0.93 |
|  | 578.53_29.19 | 0.91 |
|  | 577.52_29.19 | 0.90 |
|  | 859.78_29.17 | 0.88 |
|  | 638.57_19.38 / DG (36:2) | 0.88 |
|  | 890.45_29.19 | 0.87 |
|  | 579.54_29.19 | 0.87 |
|  | 878.82_29.2 / TG (52:1) | 0.85 |
|  | 639.58_19.4 | 0.85 |
| green | 688.6_24.91 / ChE (20:5) | 0.85 |
|  | 971.8_26.63 | 0.85 |
|  | 689.61_24.91 | 0.84 |
|  | 949.81_27.97 | 0.85 |
|  | 303.24_24.9 | 0.85 |
|  | 970.79_26.62 / TG (60:11) | 0.84 |
|  | 966.76_25.04 | 0.84 |
|  | 672.58_24.9 | 0.84 |
|  | 969.78_25.85 | 0.84 |
|  | 690.61_24.91 | 0.83 |
| midnightblue | 398.4_27.8 | 0.91 |
|  | 397.39_27.8 | 0.91 |
|  | 398.4_29.27 | 0.90 |
|  | 397.39_29.28 | 0.90 |
|  | 681.64_26.99 / ChE (20:0) | 0.89 |
|  | 680.64_27.01 | 0.89 |
|  | 682.65_26.98 | 0.89 |
|  | 384.38_27 | 0.88 |
|  | 399.4_27.8 | 0.87 |
|  | 383.37_27.01 | 0.87 |
| greenyellow | 670.65_28.82 | 0.76 |
|  | 370.36_28.83 | 0.75 |
|  | 632.64_21.49 | 0.75 |
|  | 671.66_28.82 | 0.74 |
|  | 369.36_28.96 | 0.74 |
|  | 619.63_20.87 | 0.73 |
|  | 633.64_21.48 | 0.72 |
|  | 618.62_20.87 | 0.72 |
|  | 651.65_21.51 | 0.71 |
|  | 636.63_20.87 / Cer(d41:1) | 0.70 |
| orange | 795.32_18.56 | 0.72 |
|  | 773.31_16.99 | 0.70 |
|  | 794.94_18.55 | 0.68 |
|  | 183.51_16.97 | 0.68 |
|  | 762.06_16.99 | 0.68 |
|  | 761.04_16.98 | 0.63 |
|  | 630.62_20.45 | 0.58 |
|  | 631.62_20.46 | 0.57 |
|  | 791.63_18.57 | 0.57 |
|  | 634.65_21.53 / Cer(d42:0) | 0.57 |

kME represents the module membership value for individual protein in assigned module.

**Table S3. Top 10 drivers in five selected protein modules.**

| Module | Uniprot ID | Abbreviations | Protein | kME |
| --- | --- | --- | --- | --- |
| lightgreen | P05186 | Alkaline phosphatase, bone | Alkaline phosphatase | 0.81 |
|  | P17213 | BPI | Bactericidal permeability-increasing protein | 0.79 |
|  | P02788 | Lactoferrin | Lactotransferrin | 0.76 |
|  | P78380 | LOX-1 | Oxidized low-density lipoprotein receptor 1 | 0.70 |
|  | P14780 | MMP-9 | Matrix metalloproteinase-9 | 0.69 |
|  | P16403 | Histone H1.2 | Histone H1.2 | 0.65 |
|  | O75594 | PGRP-S | Peptidoglycan recognition protein 1 | 0.50 |
|  | P07451 | Carbonic.anhydrase.III | Carbonic anhydrase 3 | 0.29 |
|  | P02649 | Apo E4 | Apolipoprotein E ε4 | 0.22 |
|  | P02649 | Apo E3 | Apolipoprotein E ε3 | 0.20 |
| lightcyan | Q96KQ7 | HMTase G9a | Histone-lysine N-methyltransferase EHMT2 | 0.65 |
|  | P01282 | Vasoactive Intestinal Peptide | Vasoactive.Intestinal.Peptides | 0.61 |
|  | P20783 | Neurotrophin-3 | Neurotrophin-3 | 0.58 |
|  | P00734 | Thrombin | Prothrombin | 0.56 |
|  | P01031 | C5 | Complement C5 | 0.53 |
|  | Q96P31 | FCRL3 | Fc receptor-like protein 3 | 0.53 |
|  | P22079 | Lactoperoxidase | Lactoperoxidase | 0.52 |
|  | P01031 | C5b,6 Complex | Complement C5b-C6 | 0.50 |
|  | P22223 | P-Cadherin | Cadherin-3 | 0.50 |
|  | P16860 | BNP-32 | Natriuretic peptides B | 0.50 |
| cyan | O95243 | MBD4 | Methyl-CpG-binding domain protein 4 | 0.66 |
|  | Q969D9 | TSLP | Thymic stromal lymphopoietin | 0.56 |
|  | Q9UHX3 | EMR2 | Adhesion G protein-coupled receptor E2 | 0.52 |
|  | P43166 | Carbonic.anhydrase.VII | Carbonic anhydrase 7 | 0.52 |
|  | P08311 | Cathepsin.G | Cathepsin G | 0.51 |
|  | P03372 | ER | Estrogen receptor | 0.51 |
|  | Q9HC73 | TSLP R | Cytokine receptor-like factor 2 | 0.49 |
|  | P09758 | GA733-1 protein | Tumor-associated calcium signal transducer 2 | 0.46 |
|  | P28908 | CD30 | Tumor necrosis factor receptor superfamily member 8 | 0.41 |
|  | Q9H293 | IL-17E | Interleukin-25 | 0.31 |
| red | P19438 | TNF sR-I | Tumor necrosis factor receptor superfamily member 1A | 0.74 |
|  | Q6NW40 | RGM-B | RGM domain family member B | 0.73 |
|  | P00746 | Factor D | Complement factor D | 0.69 |
|  | P03973 | SLPI | Antileukoproteinase | 0.64 |
|  | Q16627 | HCC-1 | C-C motif chemokine 14 | 0.60 |
|  | P61626 | Lysozyme | Lysozyme C | 0.58 |
|  | P80188 | Lipocalin 2 | Neutrophil gelatinase-associated lipocalin | 0.58 |
|  | Q03405 | suPAR | Urokinase plasminogen activator surface receptor | 0.58 |
|  | P30040 | ERp29 | Endoplasmic reticulum resident protein 29 | 0.57 |
|  | Q16663 | MIP-5 | C-C motif chemokine 15 | 0.56 |
| yellow | Q13478 | IL-18 Ra | Interleukin-18 receptor 1 | 0.64 |
|  | Q9Y286 | Siglec-7 | Sialic acid-binding Ig-like lectin 7 | 0.55 |
|  | P21810 | Biglycan | Biglycan | 0.53 |
|  | O60486 | Plexin C1 | Plexin-C1 | 0.48 |
|  | Q03154 | Aminoacylase-1 | Aminoacylase-1 | 0.47 |
|  | P05156 | Factor I | Complement factor I | 0.46 |
|  | P00797 | Renin | Renin | 0.44 |
|  | P01833 | pIgR | Polymeric immunoglobulin receptor | 0.43 |
|  | Q6UWV6 | ENPP7 | Ectonucleotide pyrophosphatase/phosphodiesterase family member 7 | 0.42 |
|  | P12830 | Cadherin-1 | Cadherin-1 | 0.41 |

kME represents the module membership value for individual protein in assigned module.

**Table S4. Summary of gene set enrichment analyses and gene ontology enrichment analysis of protein modules.**

| **Modules** | **Biological process** | **Cellular component** | **Molecular function** | **KEGG pathway** | **Reactome pathway** |
| --- | --- | --- | --- | --- | --- |
| **lightgreen** | Neutrophil mediated immunity | Vesicle lumen | Peptidase regulator activity | Cytokine-cytokine receptor interaction | Neutrophil degranulation |
|  | Granulocyte activation | Specific granule | Glycosaminoglycan binding | Osteoclast differentiation | Antimicrobial peptides |
|  | Multi-organism cellular process | Tertiary granule | Immunoglobulin binding | Phagosome | Innate immune system |
|  | STAT cascade | Blood microparticle | Lipopolysaccharide binding | JAK-STAT signaling pathway | Immune System |
|  | Negative regulation of response to external stimulus | Extracellular matrix | Proteoglycan binding | Thiamine metabolism | Binding and uptake of ligands by scavenger receptors |
|  | Biomineral tissue development | Endocytic vesicle | Enzyme inhibitor activity | Nitrogen metabolism | Chylomicron clearance |
|  | Modification of morphology or physiology of other organism | Receptor complex | Sulfur compound binding | Transcriptional misregulation in cancer | Post-translational modification: synthesis of gpi-anchored proteins |
|  | Cell killing | Platelet alpha granule | Transmembrane receptor protein kinase activity | Pathways in cancer | Metal sequestration by antimicrobial proteins |
|  | Negative regulation of proteolysis | Anchored component of membrane | Cytokine binding | Folate biosynthesis | Post-translational protein phosphorylation |
|  | Regulation of inflammatory response | Endolysosome | Protein tyrosine kinase activity | Thyroid cancer | Chylomicron remodeling |
|  | Ossification |  |  |  |  |
|  | Peptidyl-tyrosine modification |  |  |  |  |
|  | Regulation of peptidase activity |  |  |  |  |
|  | Negative regulation of hydrolase activity |  |  |  |  |
|  | Humoral immune response |  |  |  |  |
|  | Leukocyte differentiation |  |  |  |  |
|  | Regulation of protein serine/threonine kinase activity |  |  |  |  |
|  | Tumor necrosis factor superfamily cytokine production |  |  |  |  |
|  | Tissue remodeling |  |  |  |  |
|  | Regulation of innate immune response |  |  |  |  |
|  | Ameboidal-type cell migration |  |  |  |  |
|  | Cytolysis |  |  |  |  |
|  | Negative regulation of cell activation |  |  |  |  |
|  | Leukocyte migration |  |  |  |  |
| **cyan** | Positive regulation of cytokine production | Vesicle lumen | Receptor ligand activity | Cytokine-cytokine receptor interaction | Immune system |
|  | Mast cell activation | Platelet alpha granule | Serine hydrolase activity | JAK-STAT signaling pathway | Signaling by interleukins |
|  | Tumor necrosis factor superfamily cytokine production | Specific granule | Endopeptidase activity | Complement and coagulation cascades | Cytokine signaling in immune system |
|  | Interleukin-5 production | Receptor complex | Cytokine binding | Pathways in cancer | Interleukin-4 and interleukin-13 signaling |
|  | Protein maturation | Side of membrane | Cytokine receptor binding | Adipocytokine signaling pathway | Terminal pathway of complement |
|  | Leukocyte activation involved in inflammatory response | Extracellular matrix | Growth factor receptor binding | Il-17 signaling pathway | Regulation of complement cascade |
|  | Regulation of inflammatory response | Blood microparticle | Phosphatidylinositol bisphosphate kinase activity | Amoebiasis | Complement cascade |
|  | Regulation of immune effector process | Apical part of cell | Phosphatidylinositol 3-kinase activity | Hematopoietic cell lineage | Innate immune system |
|  | Peptidyl-tyrosine modification | Tertiary granule | Cytokine receptor activity | Endocrine resistance | Activation of matrix metalloproteinases |
|  | Humoral immune response | Perinuclear endoplasmic reticulum | Guanyl-nucleotide exchange factor activity | Nitrogen metabolism | Mapk1/mapk3 signaling |
|  | Interleukin-6 production |  |  |  |  |
|  | Neuroinflammatory response |  |  |  |  |
|  | STAT cascade |  |  |  |  |
|  | Acute inflammatory response |  |  |  |  |
|  | Regulation of leukocyte activation |  |  |  |  |
|  | Positive regulation of cell activation |  |  |  |  |
|  | Reactive nitrogen species metabolic process |  |  |  |  |
|  | Macrophage activation |  |  |  |  |
|  | Cytokine secretion |  |  |  |  |
|  | Protein activation cascade |  |  |  |  |
|  | Protein kinase B signaling |  |  |  |  |
|  | Interleukin-13 production |  |  |  |  |
|  | Reactive oxygen species metabolic process |  |  |  |  |
| **lightcyan** | Protein activation cascade | Endoplasmic reticulum lumen | Receptor ligand activity | Complement and coagulation cascades | Transport of gamma-carboxylated protein precursors from the endoplasmic reticulum to the golgi apparatus |
|  | Insulin-like growth factor receptor signaling pathway | Golgi lumen | Cytokine receptor binding | P53 signaling pathway | Gamma-carboxylation of protein precursors |
|  | Epithelial cell proliferation | Protein-lipid complex | Phospholipid binding | Longevity regulating pathway | Removal of aminoterminal propeptides from gamma-carboxylated proteins |
|  | Regulation of chemotaxis | Platelet alpha granule | Insulin receptor binding | Vitamin digestion and absorption | Gamma-carboxylation, transport, and amino-terminal cleavage of proteins |
|  | Leukocyte migration | Blood microparticle | Hormone receptor binding | Glycosphingolipid biosynthesis | Regulation of insulin-like growth factor (igf) transport and uptake by insulin-like growth factor binding proteins (igfbps) |
|  | Regulation of plasma lipoprotein particle levels | Vesicle lumen | Cell adhesion molecule binding | Transcriptional misregulation in cancer | Formation of fibrin clot (clotting cascade) |
|  | Cell growth | Endoplasmic reticulum exit site | Fibronectin binding | Prion diseases | Gamma carboxylation, hypusine formation and arylsulfatase activation |
|  | Regulation of body fluid levels | Smooth endoplasmic reticulum | Lipoprotein particle receptor binding | Aldosterone-regulated sodium reabsorption | Regulation of complement cascade |
|  | Positive regulation of response to external stimulus | Endosome lumen | Serine hydrolase activity | Fat digestion and absorption | Complement cascade |
|  | Protein-containing complex remodeling | Extrinsic component of membrane | Peptidase regulator activity | Ras signaling pathway | Synthesis, secretion, and deacylation of ghrelin |
| **red** | Humoral immune response | Vesicle lumen | Receptor ligand activity | Cytokine-cytokine receptor interaction | Immune system |
|  | Response to tumor necrosis factor | Extracellular matrix | Serine hydrolase activity | Mapk signaling pathway | Regulation of insulin-like growth factor (igf) transport and uptake by insulin-like growth factor binding proteins (igfbps) |
|  | Leukocyte migration | Specific granule | Endopeptidase activity | Rap1 signaling pathway | Signaling by interleukins |
|  | Cell chemotaxis | Endoplasmic reticulum lumen | Protein tyrosine kinase activity | Fluid shear stress and atherosclerosis | Cytokine signaling in immune system |
|  | Response to interleukin-1 | Vacuolar lumen | Phosphatidylinositol bisphosphate kinase activity | Nf-kappa b signaling pathway | Neutrophil degranulation |
|  | Neutrophil mediated immunity | Primary lysosome | Glycosaminoglycan binding | Hif-1 signaling pathway | Activation of matrix metalloproteinases |
|  | Granulocyte activation | Anchored component of membrane | Phosphatidylinositol 3-kinase activity | Tuberculosis | Innate immune system |
|  | Positive chemotaxis | Extrinsic component of membrane | Enzyme inhibitor activity | Kaposi sarcoma-associated herpesvirus infection | Degradation of the extracellular matrix |
|  | Positive regulation of response to external stimulus | Ficolin-1-rich granule | Cytokine receptor binding | Chemokine signaling pathway | Interleukin-10 signaling |
|  | Regulation of vasculature development | Pigment granule | Peptidase regulator activity | Osteoclast differentiation | Post-translational protein phosphorylation |
|  | Negative regulation of hydrolase activity |  |  |  |  |
|  | Negative regulation of proteolysis |  |  |  |  |
|  | Defense response to other organism |  |  |  |  |
|  | Angiogenesis |  |  |  |  |
|  | Adaptive immune response |  |  |  |  |
|  | Peptidyl-tyrosine modification |  |  |  |  |
|  | Extracellular structure organization |  |  |  |  |
|  | ERK1 and ERK2 cascade |  |  |  |  |
|  | Production of molecular mediator of immune response |  |  |  |  |
|  | Response to interferon-gamma |  |  |  |  |
|  | Regulation of peptidase activity |  |  |  |  |
|  | Response to mechanical stimulus |  |  |  |  |
|  | Platelet degranulation |  |  |  |  |
|  | Multi-multicellular organism process |  |  |  |  |
|  | Transmembrane receptor protein serine/threonine kinase signaling pathway |  |  |  |  |
|  | T cell activation |  |  |  |  |
|  | Response to oxygen levels |  |  |  |  |
| **yellow** | Humoral immune response | Extracellular matrix | Serine hydrolase activity | Staphylococcus aureus infection | Platelet degranulation |
|  | Protein activation cascade | Vesicle lumen | Endopeptidase activity | Cytokine-cytokine receptor interaction | Response to elevated platelet cytosolic ca2+ |
|  | Platelet degranulation | Blood microparticle | Receptor ligand activity | Phagosome | Immune system |
|  | Adaptive immune response | Platelet alpha granule | Immunoglobulin binding | Complement and coagulation cascades | Extracellular matrix organization |
|  | Extracellular structure organization | Vacuolar lumen | Carbohydrate binding | Tuberculosis | Complement cascade |
|  | ERK1 and ERK2 cascade | Side of membrane | Proteoglycan binding | Malaria | Innate Immune System |
|  | Acute inflammatory response | Receptor complex | Transmembrane receptor protein kinase activity | Tgf-beta signaling pathway | Initial triggering of complement |
|  | Regulation of immune effector process | Endoplasmic reticulum lumen | Cytokine receptor binding | Intestinal immune network for iga production | Platelet activation, signaling and aggregation |
|  | Negative regulation of cell adhesion | Endolysosome | Protein tyrosine kinase activity | Leishmaniasis | Formation of fibrin clot (clotting cascade) |
|  | Granulocyte activation | Primary lysosome | Growth factor binding | Egfr tyrosine kinase inhibitor resistance | Neutrophil degranulation |
|  | Regulation of inflammatory response |  | Peptidase regulator activity |  |  |
|  | Lymphocyte mediated immunity |  | Cytokine receptor activity |  |  |
|  | Neutrophil mediated immunity |  | Extracellular matrix structural constituent |  |  |
|  | Negative regulation of proteolysis |  |  |  |  |
|  | Collagen metabolic process |  |  |  |  |
|  | Regulation of vesicle-mediated transport |  |  |  |  |
|  | Leukocyte cell-cell adhesion |  |  |  |  |
|  | Protein maturation |  |  |  |  |
|  | Regulation of peptidase activity |  |  |  |  |
|  | Negative regulation of hydrolase activity |  |  |  |  |
|  | Connective tissue development |  |  |  |  |
|  | Defense response to other organism |  |  |  |  |
|  | Receptor-mediated endocytosis |  |  |  |  |
|  | Regulation of cell-cell adhesion |  |  |  |  |
|  | Cell recognition |  |  |  |  |
|  | Response to interleukin-1 |  |  |  |  |
|  | Modification of morphology or physiology of other organism |  |  |  |  |
|  | Positive regulation of defense response |  |  |  |  |
|  | Cell killing |  |  |  |  |
|  | Phagocytosis |  |  |  |  |
|  | Cell-substrate adhesion |  |  |  |  |
|  | Gland development |  |  |  |  |
|  | Response to tumor necrosis factor |  |  |  |  |
|  | Epithelial cell apoptotic process |  |  |  |  |
|  | Regulation of peptide secretion |  |  |  |  |
|  | Mesenchyme development |  |  |  |  |
|  | Response to fungus |  |  |  |  |
|  | Interleukin-10 production |  |  |  |  |
|  | Protein-lipid complex subunit organization |  |  |  |  |
|  | Exocrine system development |  |  |  |  |
|  | Response to nutrient levels |  |  |  |  |
|  | Production of molecular mediator of immune response |  |  |  |  |
|  | Response to interferon-gamma |  |  |  |  |
|  | Peptidyl-tyrosine modification |  |  |  |  |
|  | Positive regulation of cell adhesion |  |  |  |  |
|  | Heterotypic cell-cell adhesion |  |  |  |  |
|  | Positive regulation of secretion |  |  |  |  |
|  | Odontogenesis |  |  |  |  |

GO terms and pathways do not pass Benjamini-Hochberg FDR correction were highlighted in tan.

**Table S5. Correlations between lipid modules/protein modules and AD genetic variants.**

| Lipid modules | Locus | Variant | chromosome | *p* value |
| --- | --- | --- | --- | --- |
| green | MEF2C | rs190982 | 5 | 4.75e-03 |
|  | SORL1 | rs79398263 | 7 | 2.99e-02 |
|  | ABCA7 | rs3752246 | 19 | 3.80e-02 |
| darkturquoise | IL-34 | rs4985556 | 16 | 1.20e-03 |
|  | MEF2C | rs190982 | 5 | 5.50e-03 |
|  | CR1 | rs4844610 | 1 | 1.23e-02 |
|  | SORL1 | rs79398263 | 7 | 1.45e-02 |
| midnightblue | FERMT2 | rs17125924 | 14 | 4.69e-02 |
| Protein modules | **Locus** | **Variant** | **chromosome** | ***p* value** |
| lightgreen | IQCK | rs7185636 | 16 | 2.48e-02 |
|  | PTK2B | rs73223431 | 8 | 2.50e-02 |
| red | MS4 | rs7933202 | 11 | 7.90e-03 |
|  | HLA -DRB1 | rs9271058 | 6 | 3.30e-02 |
| lightcyan | CASS4 | rs6024870 | 20 | 2.60e-03 |
|  | APOE | rs429358 | 19 | 1.27e-02 |
|  | HLA -DRB1 | rs9271058 | 6 | 1.64e-02 |
| cyan | NDUFAF6 | rs4735340 | 8 | 4.69e-03 |
|  | IL-34 | rs4985556 | 16 | 1.35e-02 |
|  | PICALM | rs3851179 | 11 | 4.01e-02 |
|  | APOE | rs429358 | 19 | 4.8e-02 |

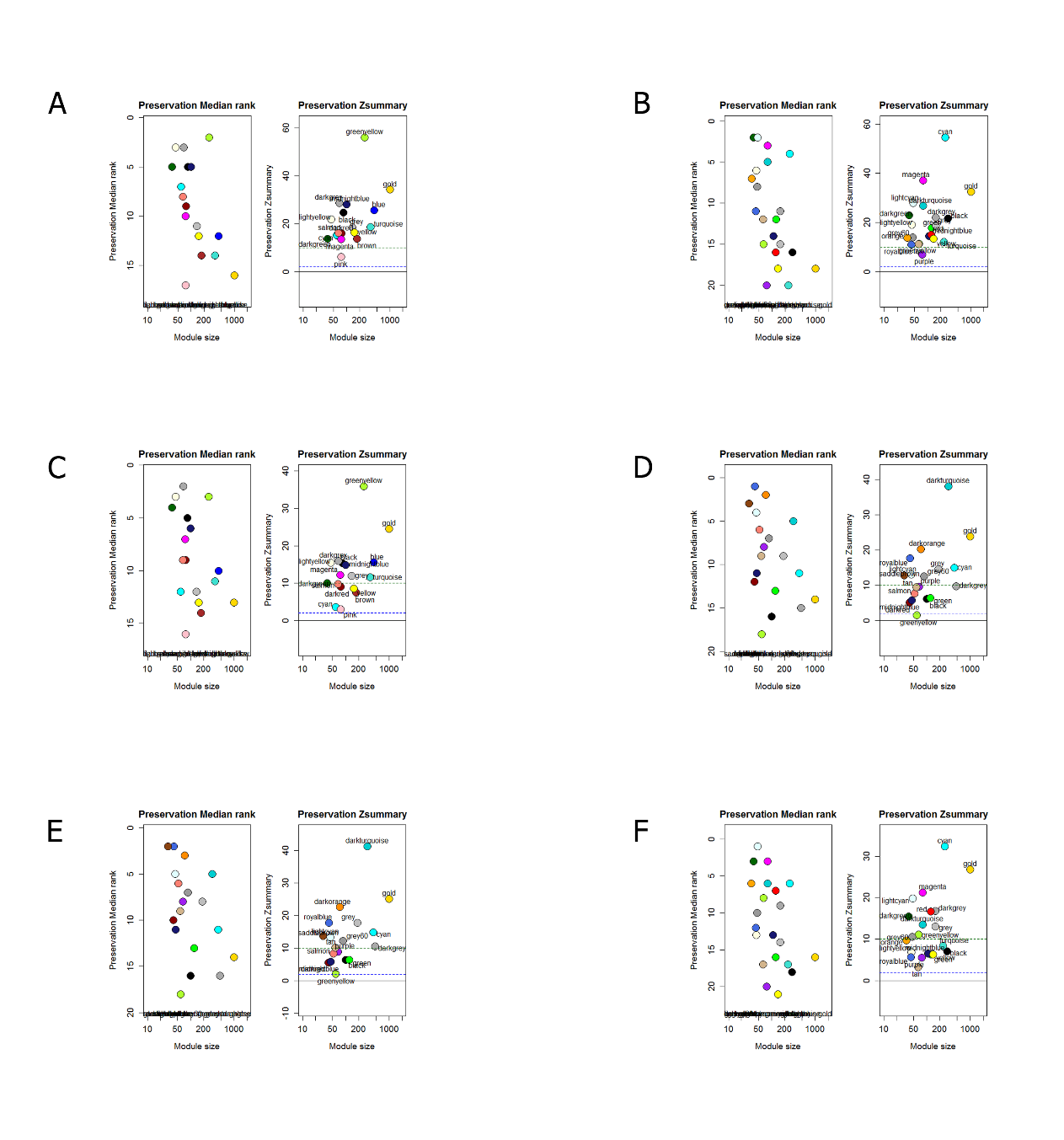

**Figure S1. Preservation summary plots for AD, MCI and control sub datasets in lipidomics dataset.** A. AD sub dataset (test) and control sub dataset (reference); B. control sub dataset (test) and AD sub dataset (reference); C. AD sub dataset (test) and MCI sub dataset (reference); D. MCI sub dataset (test) and AD sub dataset (reference); E. MCI sub dataset (test) and control sub dataset (reference); F. control sub dataset (test) and MCI sub dataset (reference).

Z*_summary_* is the summary preservation statistics. Y-axis represents preservation statistics for the corresponding module in the case data sets, and x-axis is the lipid feature numbers in each module. The dashed blue and green lines indicate the thresholds Z=2 and Z=10, respectively. Z*_summary_* < 2 implies no evidence for module preservation, 2 < Z*_summary_* < 10 implies weak to moderate evidence, and Z*_summary_* > 10 implies strong evidence for module preservation.

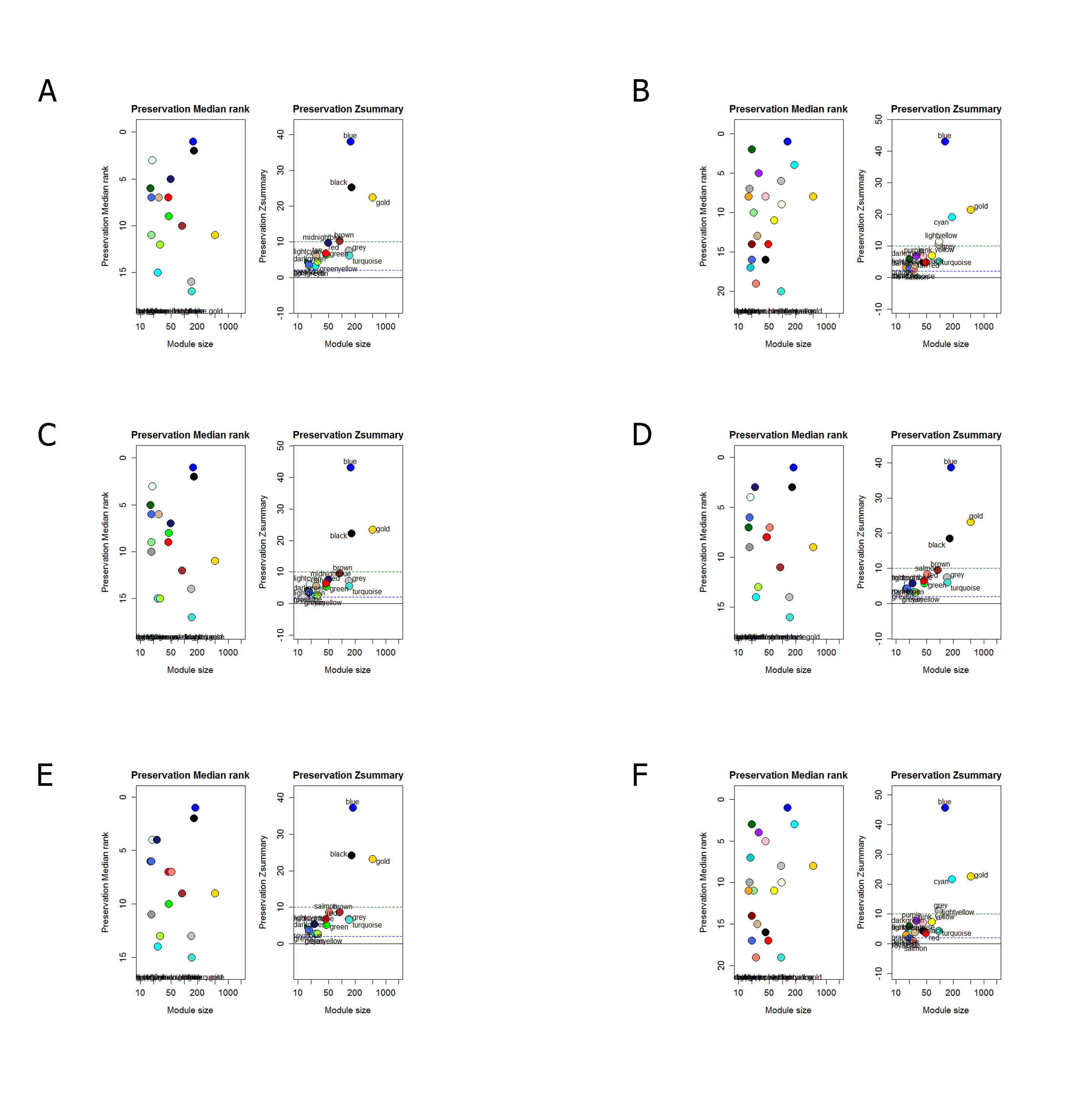

**Figure S2.** **Preservation summary plots for AD, MCI and control sub datasets in ANM proteomics dataset.** A. AD sub dataset (test) and control sub dataset (reference); B. control sub dataset (test) and AD sub dataset (reference); C. AD sub dataset (test) and MCI sub dataset (reference); D. MCI sub dataset (test) and AD sub dataset (reference); E. MCI sub dataset (test) and control sub dataset (reference); F. control sub dataset (test) and MCI sub dataset (reference).

Z*_summary_* is the summary preservation statistics. Y-axis represents preservation statistics for the corresponding module in the case data sets, and x-axis is the lipid feature numbers in each module. The dashed blue and green lines indicate the thresholds Z=2 and Z=10, respectively. Z*_summary_* < 2 implies no evidence for module preservation, 2 < Z*_summary_* < 10 implies weak to moderate evidence, and Z*_summary_* > 10 implies strong evidence for module preservation.

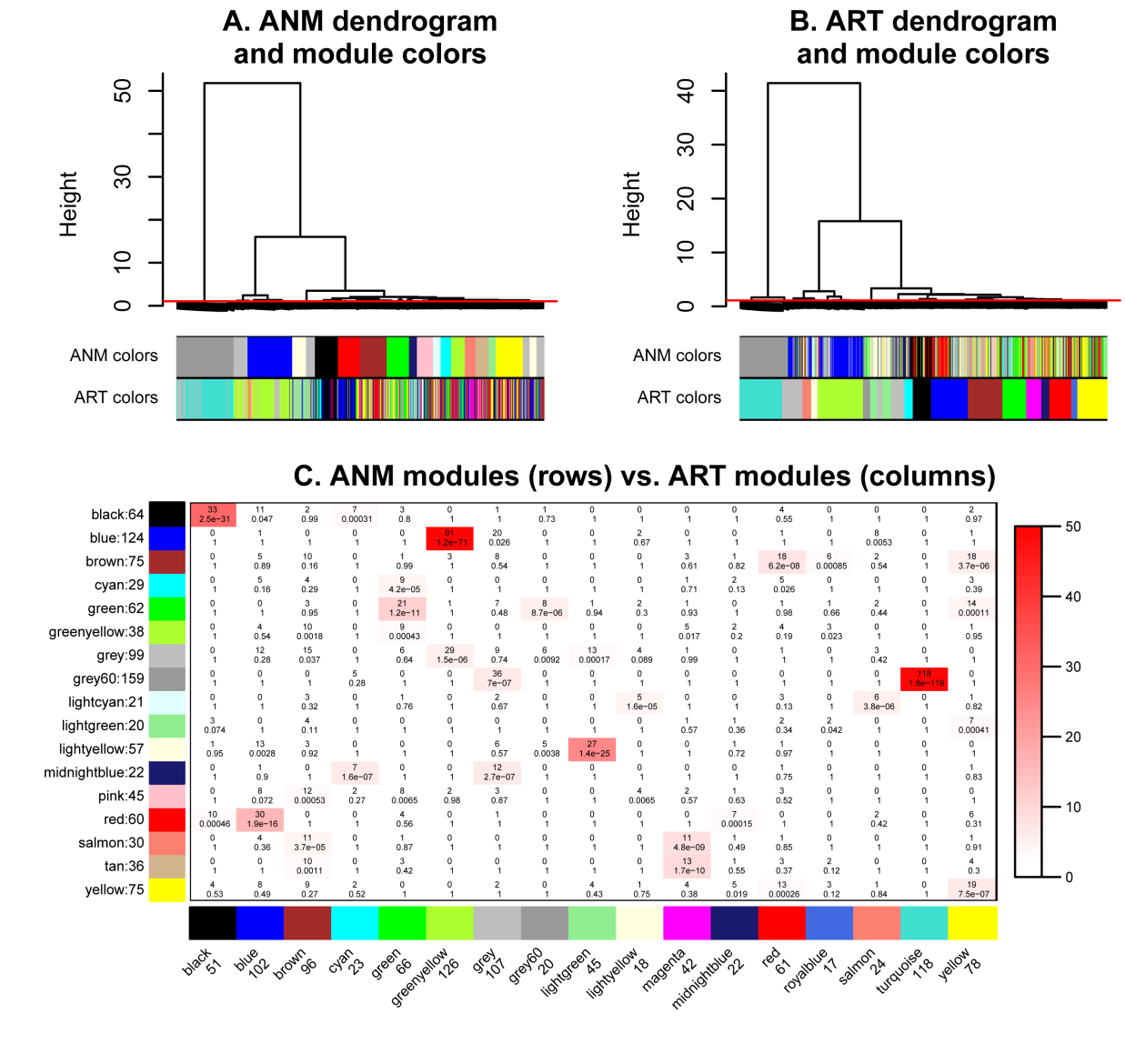

**Figure S3. Dendrogram and cross-tabulation based comparison of modules in ANM and ART protein cohort networks.** A. Hierarchical clustering tree (dendrogram) of proteins based on ANM cohort network. The color rows below the dendrogram indicate module membership in the ANM modules (defined by cutting branches of this dendrogram at the red line) and in the ART network (defined by branch cutting the dendrogram in panel B.) B. Hierarchical clustering tree of proteins based on the ART co-expression network. The color rows below the dendrogram indicate module membership in the human modules (defined by cutting branches of dendrogram in panel A.) and in the ART network (defined by branch cutting the dendrogram in this panel.) C. Cross-tabulation of ANM modules (rows) and ART modules (columns). Each row and column is labeled by the corresponding module color and the total number of proteins in the module. In the table, numbers give counts of proteins in the intersection of the corresponding row and column module. The table is color-coded by -log(p), the Fisher exact test p value, according to the color legend on the right.

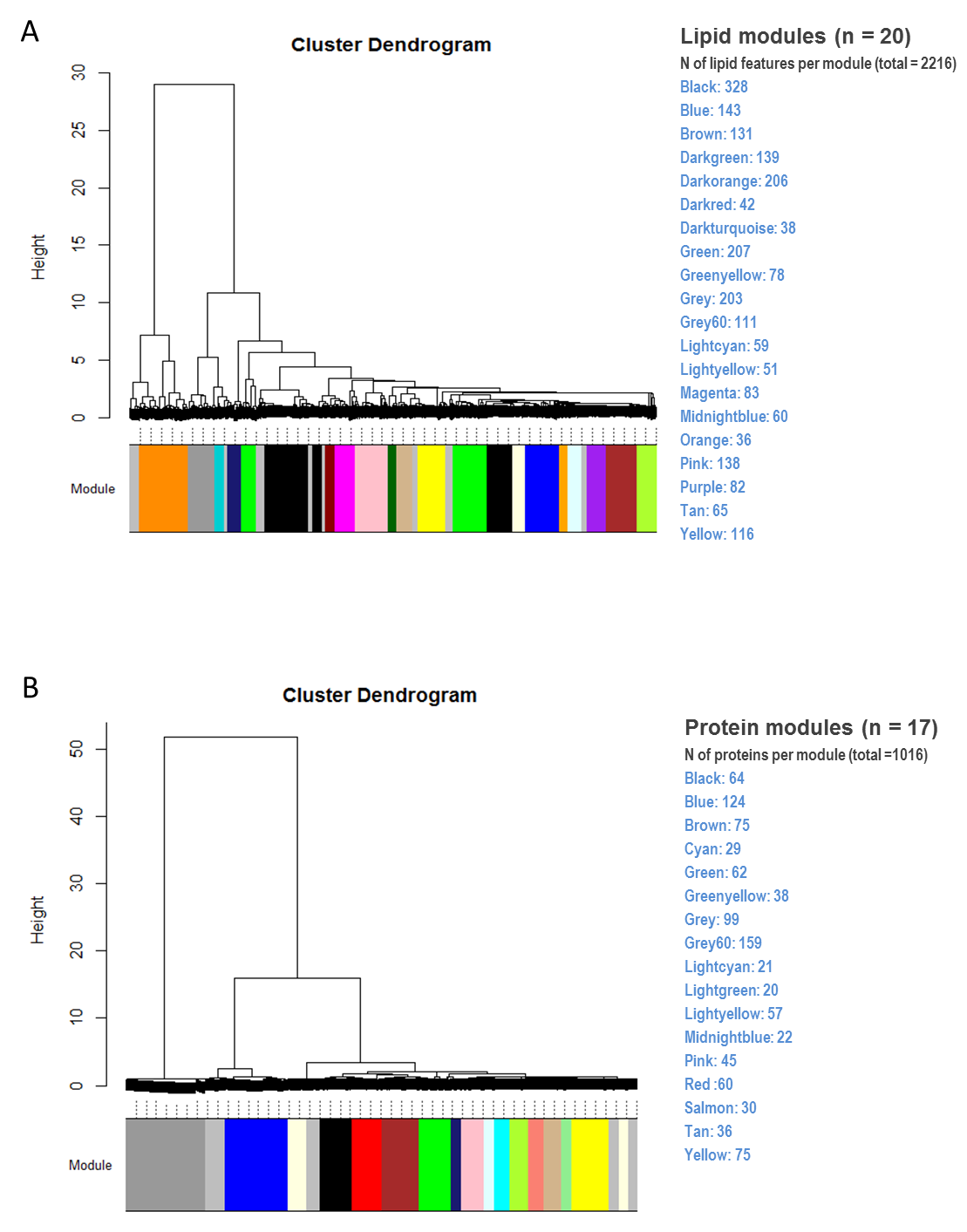

**Figure S4.** A. Cluster dendrogram of weighted lipid correlation network analysis and the number of lipid features in each module; B. Cluster dendrogram of weighted protein correlation network analysis and the number of proteins in each module.

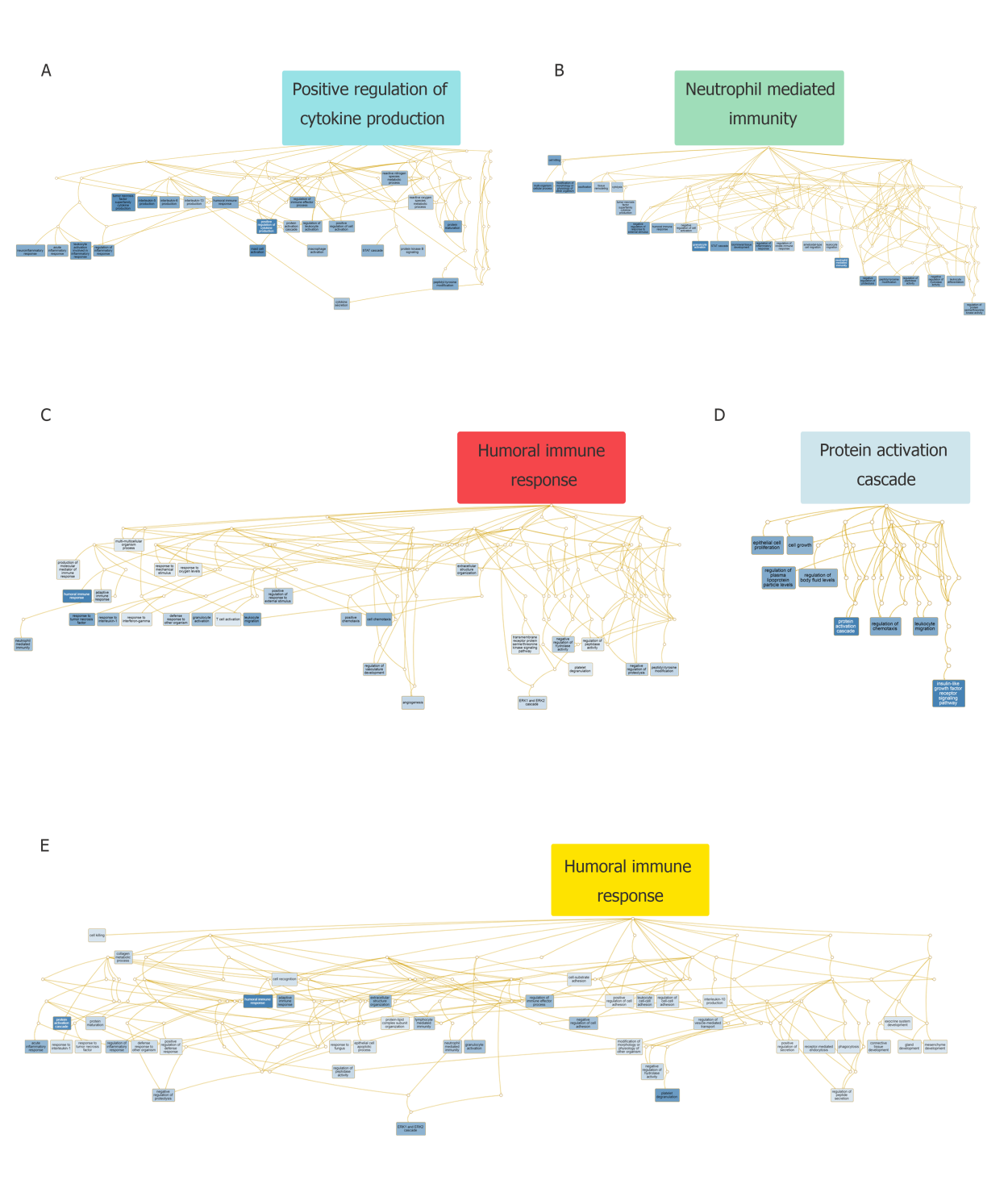

**Figure S5.** **DAGs summarize biological processes in five protein modules.** (A) cyan module; (B) lightgreen module; (C) lightcyan module; (D) red module; (E) yellow module. Each DAG includes the all biological processes passed the BH correction for each protein module. The top biological process for each module was listed in the box above the DAG, and the colour of the boxes corresponds to the colour of the module.

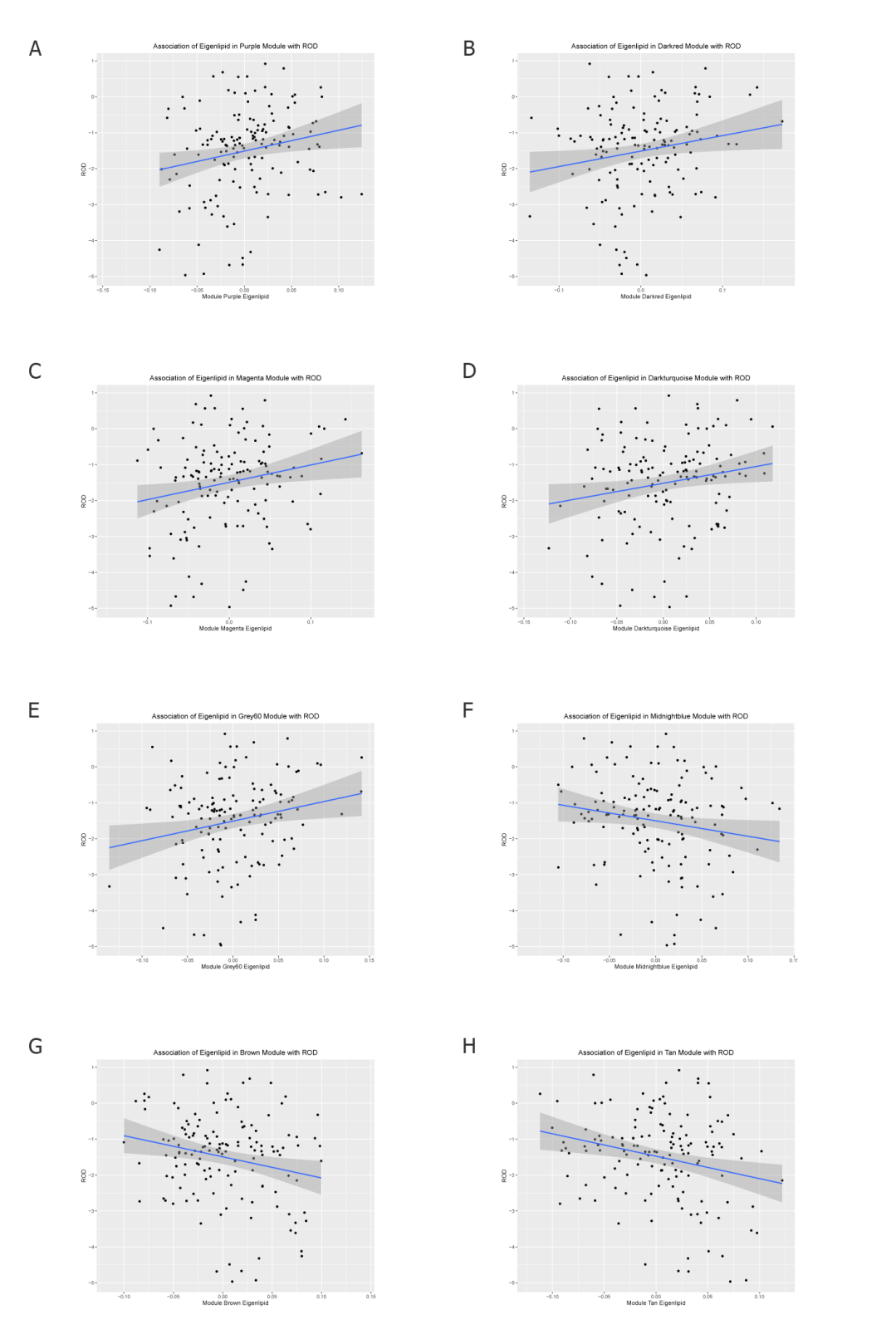

**Figure S6.** **Scatter plots of eigenlipids correlation with ROD.** (A) Eigenlipid in purple module; (B) Eigenlipid in darkred module; (C) Eigenlipid in magenta module; (D) Eigenlipid in darkturquoise module; (E) Eigenlipid in grey60 module; (F) Eigenlipid in midnightblue module; (G) Eigenlipid in brown module; (H) Eigenlipid in tan module.

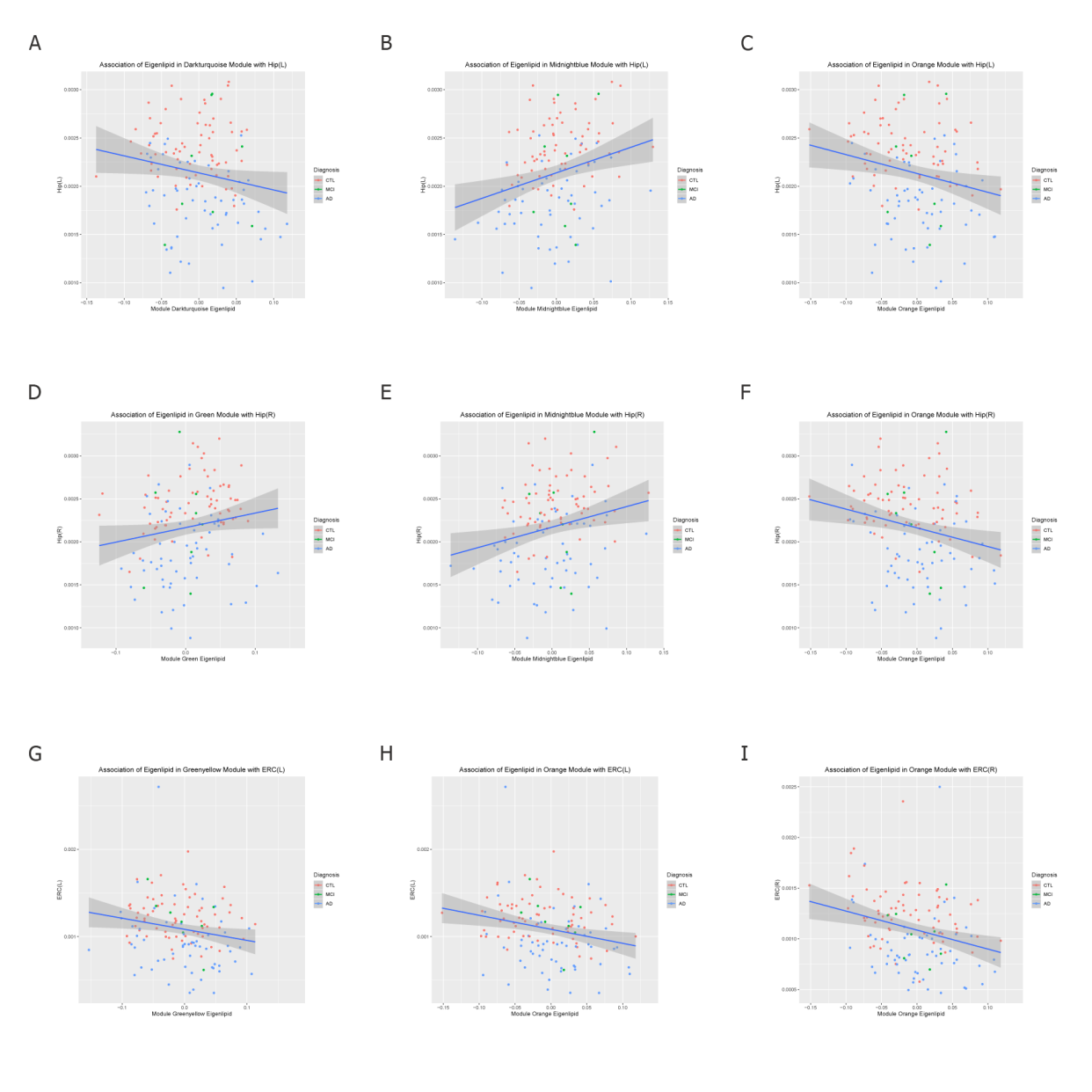

**Figure S7.** **Scatter plots of eigenlipids correlation with brain atrophy measures.** (A) Eigenlipid in darkturquoise module vs. Hippocampal left volume ; (B) Eigenlipid in midnightblue module vs. Hippocampal left volume; (C) Eigenlipid in orange module vs. Hippocampal left volume; (D) Eigenlipid in green module vs. Hippocampal right volume; (E) Eigenlipid in midnightblue module vs. Hippocampal right volume; (F) Eigenlipid in orange module vs. Hippocampal right volume; (G) Eigenlipid in greenyellow module vs. entorhinal cortex left volume; (H) Eigenlipid in orange module vs. entorhinal cortex left volume; (I) Eigenlipid in orange module vs. entorhinal cortex right volume.

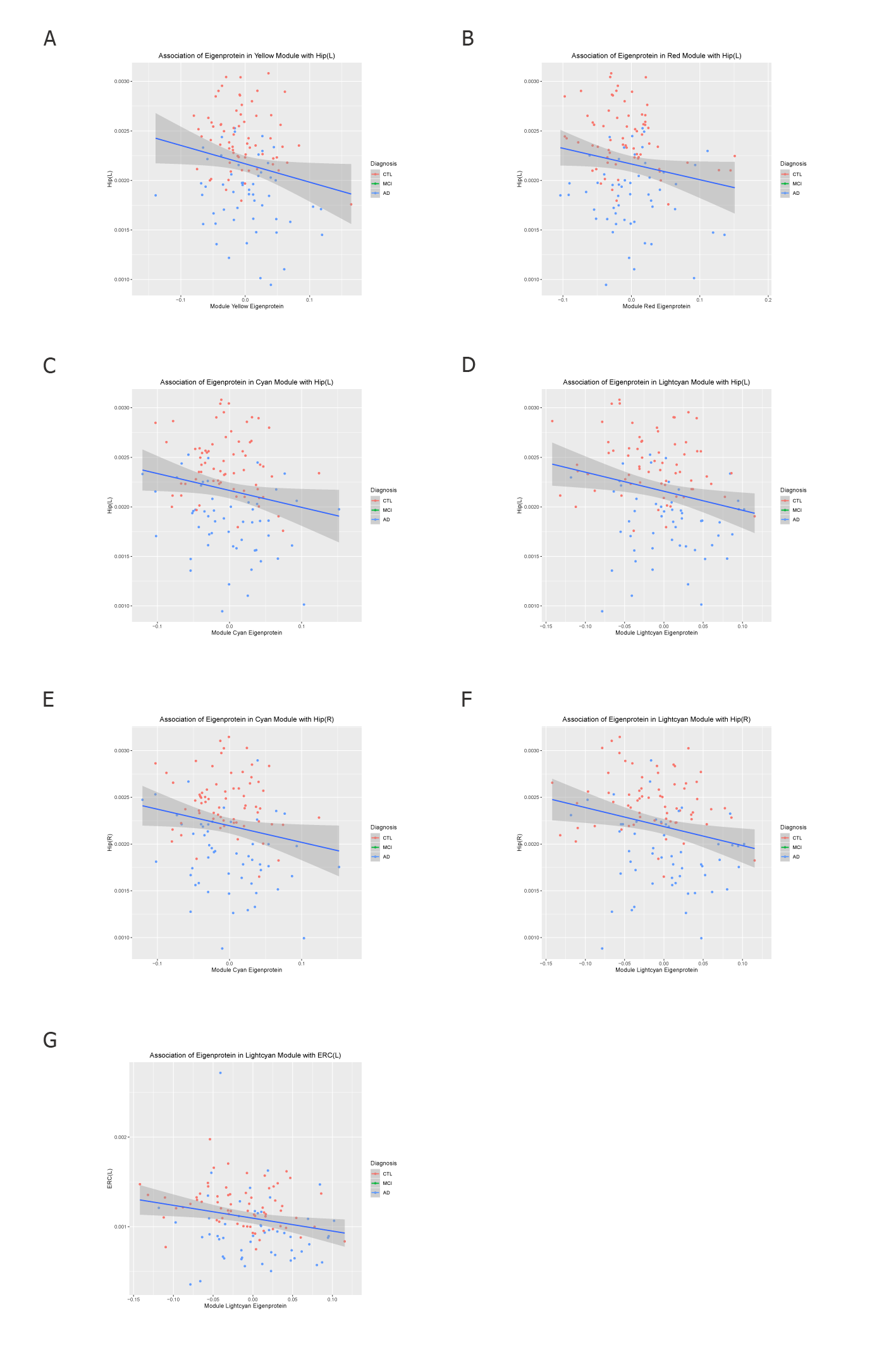

**Figure S8.** **Scatter plots of eigenproteins correlation with brain atrophy measures.** (A) Eigenprotein in yellow module vs. Hippocampal left volume ; (B) Eigenprotein in red module vs. Hippocampal left volume; (C) Eigenprotein in cyan module vs. Hippocampal left volume; (D) Eigenprotein in lightcyan module vs. Hippocampal left volume; (E) Eigenprotein in cyan module vs. Hippocampal right volume; (F) Eigenprotein in lightcyan module vs. Hippocampal right volume; (G) Eigenlipid in lightcyan module vs. entorhinal cortex left volume.

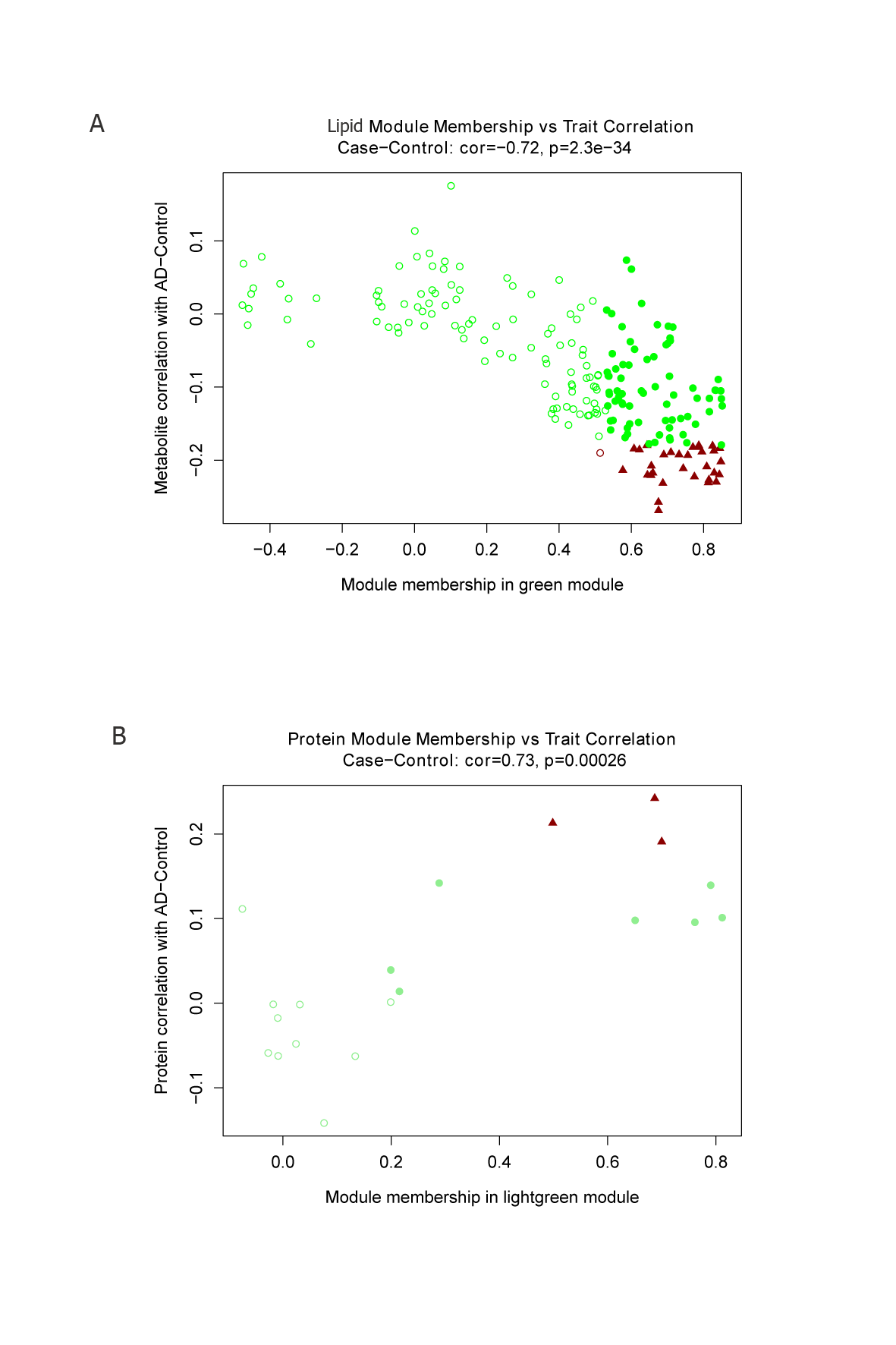

**Figure S9.** **Scatter plots of module membership versus lipids/proteins-diagnosis correlation.** (A) Lipid module membership for green module vs lipid-diagnosis correlation; (B) Protein module membership for lightgreen module vs protein-diagnosis correlation. Filled circles in represent lipids/proteins with MM > median & correlation q value (Bonferroni correction) > 0.05; filled dark red triangles represents lipid/proteins with MM > median & correlation q value (Bonferroni correction) < 0.05; hollow dark red circles represent lipid/proteins with MM < median & correlation q value (Bonferroni correction) < 0.05.

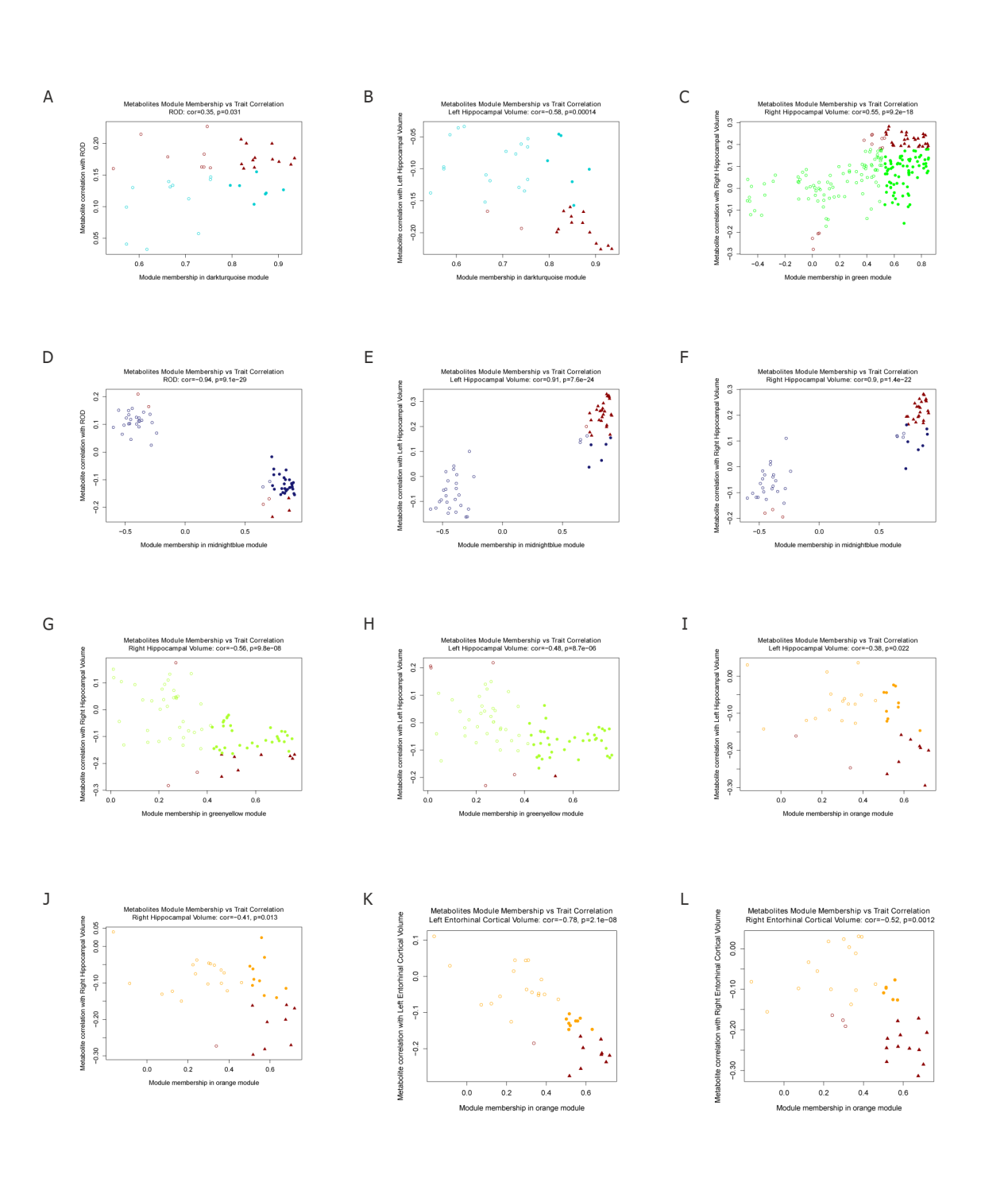

**Figure S10. Scatter plots of lipid module membership versus lipids-phenotypes correlation.** (A) kMEs in darkturquoise module vs. lipids-ROD; (B) kMEs in darkturquoise module vs. lipids-Hippocampal left volume; (C) kMEs in green module vs. lipids-Hippocampal right volume; (D) kMEs in midnightblue module vs. lipids-ROD; (E) kMEs in midnightblue module vs. lipids-Hippocampal left volume; (F) kMEs in midnightblue module vs. lipids-Hippocampal right volume; (G) kMEs in greenyellow module vs. lipids-Hippocampal right volume; (H) kMEs in greenyellow module vs. lipids-entorhinal cortex left volume; (I) kMEs in orange module vs. lipids-Hippocampal left volume; (J) kMEs in orange module vs. lipids-Hippocampal right volume; (K) kMEs in orange module vs. lipids-entorhinal cortex left volume; (L) kMEs in orange module vs. lipids-entorhinal cortex right volume.

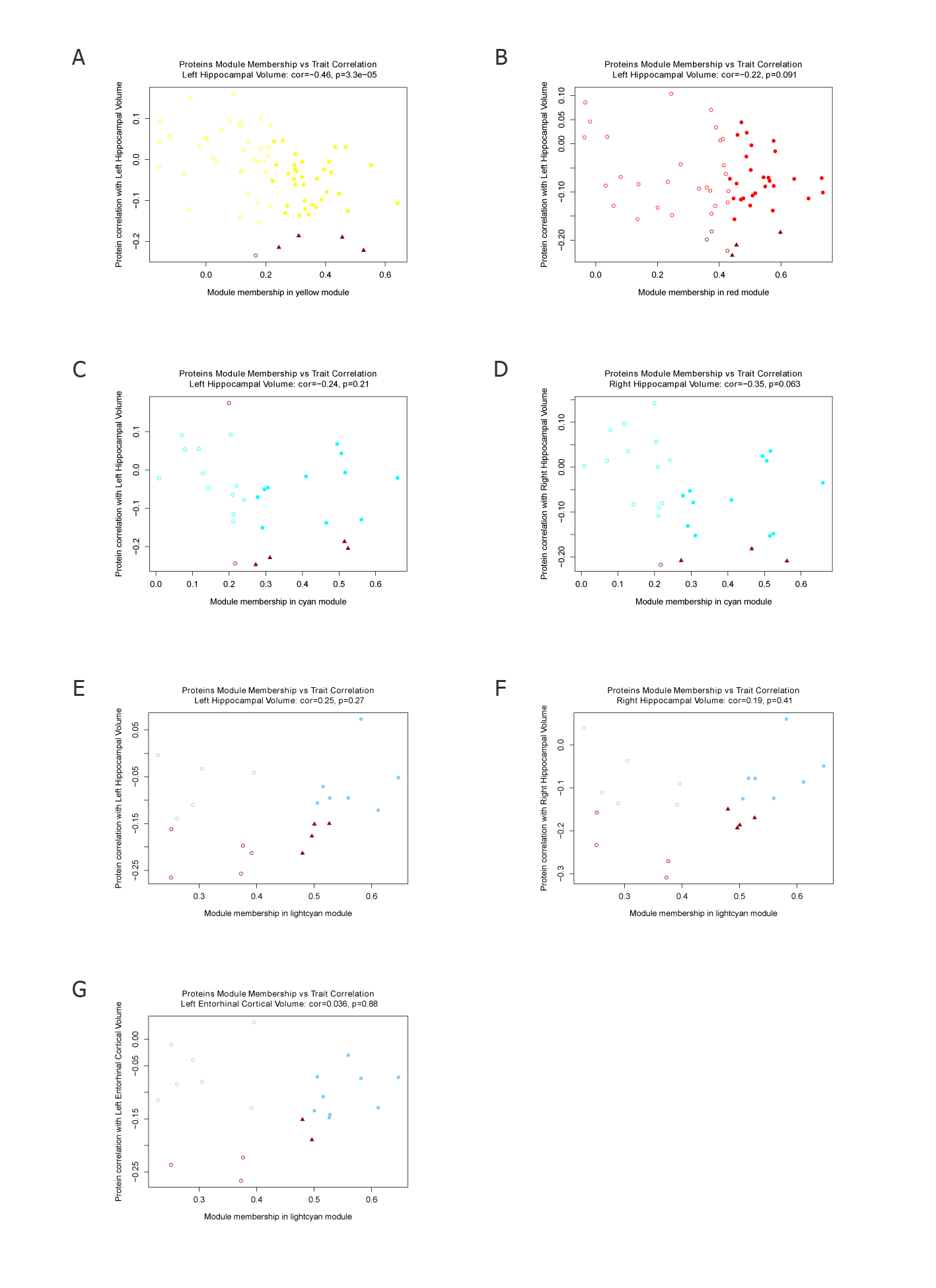

**Figure S11. Scatter plots of protein module membership versus proteins-phenotypes correlation.** (A) kMEs in yellow module vs. proteins-Hippocampal left volume; (B) kMEs in red module vs. proteins-Hippocampal left volume; (C) kMEs in cyan module vs. proteins-Hippocampal left volume; (D) kMEs in cyan module vs. proteins-Hippocampal right volume; (E) kMEs in lightcyan module vs. proteins-Hippocampal left volume; (F) kMEs in lightcyan module vs. proteins-Hippocampal right volume; (G) kMEs in lightcyan module vs. proteins-entorhinal cortex left volume.

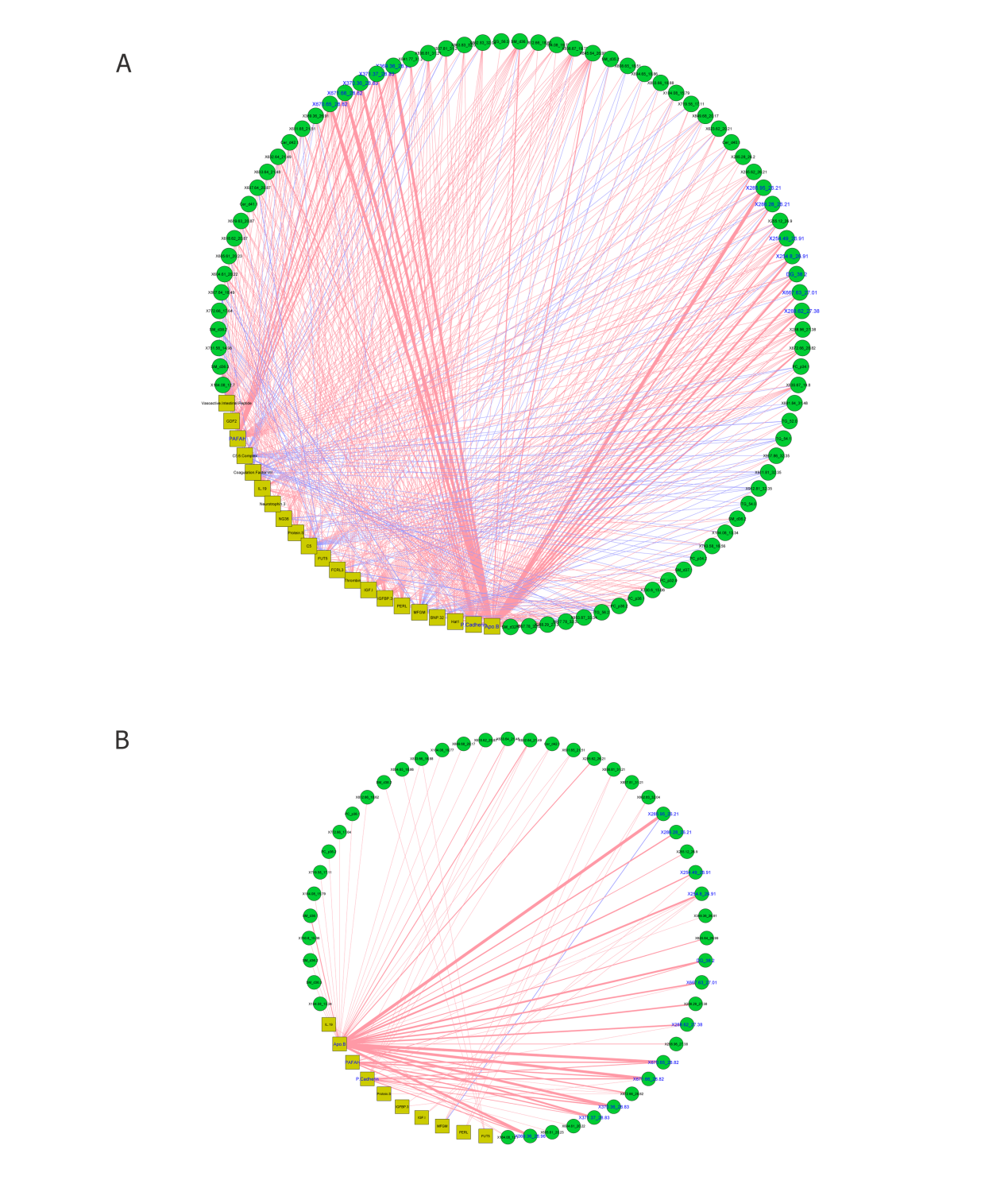

**Figure S12. Correlation networks for lipid greenyellow module and protein lightcyan module.** (A) Correlation between lipids and proteins within the range of (-1, -0.1) & (0.1, 1); (B) Correlation between lipids and proteins within the range of (-1, -0.2) & (0.2, 1). Pearson correlation was applied. Green circles represent lipids and yellow squares represent proteins. Pink edge mean positive correlation and blue edge means negative correlation. The width of the edge correlates with the strength of the correlation. Lipids/protein names were labelled in blue if correlation p values passing Holm–Bonferroni correction.

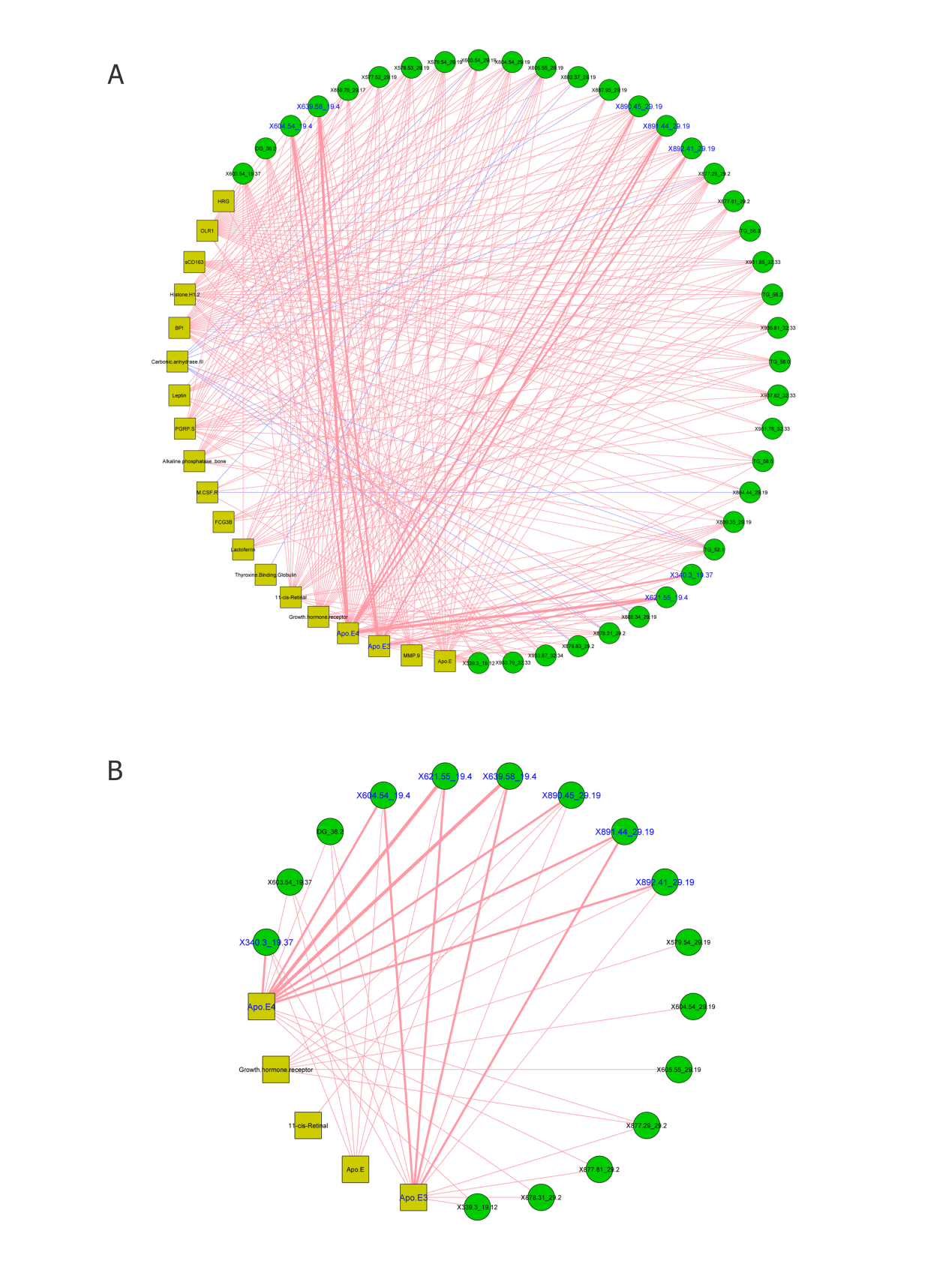

**Figure S13. Correlation networks for lipid darkturquoise module and protein lightgreen module.** (A) Correlation between lipids and proteins within the range of (-1, -0.1) & (0.1, 1); (B) Correlation between lipids and proteins within the range of (-1, -0.2) & (0.2, 1). Pearson correlation was applied. Green circles represent lipids and yellow squares represent proteins. Pink edge mean positive correlation and blue edge means negative correlation. The width of the edge correlates with the strength of the correlation. Lipids/protein names were labelled in blue if correlation p values passing Holm–Bonferroni correction.
